# *In silico* single-cell metabolism analysis unravels a new transition stage of CD8 T cells 4 days post-infection

**DOI:** 10.1101/2023.09.22.558248

**Authors:** Arpin Christophe, Franck Picard, Olivier Gandrillon

**Affiliations:** Laboratory of Biology and Modeling of the Cell, Université de Lyon, ENS de Lyon, Université Claude Bernard, CNRS UMR 5239, INSERM U1210, Lyon, France; Centre International de recherche en Infectiologie, Université de Lyon, ENS de Lyon, Université Claude Bernard, CNRS UMR 5308, INSERM U1111, Lyon, France; Inria, France

## Abstract

CD8 T cell proper differentiation during antiviral responses relies on metabolic adaptations. Herein, we investigated global metabolic activity in single CD8 T cells along an *in vivo* response by estimating metabolic fluxes from single-cell RNA-sequencing data. The approach was validated by the observation of metabolic variations known from experimental studies on global cell populations, while adding temporally detailed information and unravelling yet undescribed sections of CD8 T cell metabolism that are affected by cellular differentiation. Furthermore, inter-cellular variability in gene expression level, highlighted by single cell data, and heterogeneity of metabolic activity 4 days post-infection, revealed a new transition stage accompanied by a metabolic switch in activated cells differentiating into full-blown effectors.

## 1. INTRODUCTION

CD8 T lymphocytes are critical cytotoxic effector cells that protect against viral infections by eradicating virus-infected cells. The antiviral response typically recruits rare antigen (Ag)-specific naive CD8 T cells, resulting in their tremendous proliferation and differentiation into potent cytotoxic effectors that eliminate infected cells. After clearing of viral Ag, the majority of effector cells undergoes apoptosis leaving a small population of quiescent memory cells, which will ensure rapid and most effective secondary responses to subsequent infections with the same pathogen [1,2].

This differentiation process, which produces immediate effector and long-term protector lymphocyte populations, is accompanied by metabolic reprogramming at different stages in order to support specific bioenergetic requirements of differentiating cells [3–5]. For instance, glucose consumption by CD8 T lymphocytes relies on its cytoplasmic degradation to pyruvate that can be further catabolized to lactate, a process known as glycolysis. Alternatively, pyruvate can enter mitochondria, where its conversion into acetyl-coenzymeA (acetyl-coA) fuels the TriCarboxylic Acid cycle (TCA), which in turn activates the mitochondrial membrane Electron Transport Chain for Oxidative phosphorylation (Oxphos) of Adenosine-DiPhosphate to Adenosine-TriPhosphate. CD8 T cell can also produce energy from fatty acid oxidation (FAO) in mitochondria, which produces acetyl-coA for the TCA. The metabolism of quiescent naive CD8 T cells mostly relies on basic glycolysis and FAO [4]. Upon activation, CD8 T cells enhance glucose consumption [6] and switch to aerobic glycolysis [7,3,8,9,4,5]. This metabolic switch is mandatory for the proper differentiation of naive cells into effectors [10,11]. Later in the response, when effectors differentiate to memory cells, they undergo a new metabolic switch back to FAO [7,12,13], which is again mandatory for proper memory CD8 T cell generation [11,14,5].

Besides this well described reprogramming of energy production during CD8 T cell responses to viral infections, Ag encounter also promotes cholesterol biosynthesis [15,16] and triggers an increase in glutamine uptake and glutaminolysis [17,18], which fuels FA synthesis through α-KetoGlutarate (α-KG)-dependent citrate production [4,10]. T cell activation also redirects glyceraldehyde-3P (G3P) downstream glucose degradation towards the production of 5-phosphoribose-2P (PRPP) that fuels the pentose-phosphate pathway (PPP) to meet nucleic acid and aromatic amino acid biosynthesis demands [18]. Globally, CD8 T cell activation, proliferation and differentiation are coupled to metabolic reprogramming that supports trade-offs between energy demand and biomolecule synthesis and participates in cell differentiation. Both cell differentiation and metabolic switches are intertwined and co-regulated epigenetically [19,9,20,4,5].

These metabolic adaptations to viral challenge have been studied on global populations of CD8 T cell responders. However, effector and memory CD8 T cell populations are heterogeneous [2,21,22] and different subsets shall undergo and require specific metabolic programs [23,24]. Thus, an analysis of the metabolism of CD8 T cell responding to a viral infection at the single-cell level would be much more biologically relevant [25–27]. However, single-cell metabolomic techniques still suffer from relatively low throughput and sensitivity [28,29]. Thus, to generate a global description of the metabolism of CD8 T cell responding to a viral infection at the single-cell level, we made use of the recently described single-cell Flux Estimation Analysis (scFEA) algorithm [30] that estimates metabolite fluxes in cells from single-cell RNA-sequencing (scRNA-seq) expression data. Our results evidenced metabolic switches previously described, thus validating the approach, and revealed new metabolic perturbations that may deserve further detailed experimental validation.

Furthermore, this single-cell level analysis revealed time-dependent variation in inter-cellular heterogeneity. Indeed, we show that, as in numerous other cell differentiation systems [31], metabolic genes are subjected to a transient rise in gene expression variability allowing cells to explore the gene expression space, before selection of the most appropriate gene regulatory network (GRN) state resulting in a more homogeneous gene expression pattern in the emerging differentiated populations [32]. Interestingly, two distinct surges in metabolic gene expression variability were observed: immediately after activation and around 4 days post-infection (dpi). The genes concerned by these perturbations are involved in largely non-overlapping pans of cell metabolism, suggesting a yet unknown metabolic switch at 4 dpi, when activated cells fully commit to effector differentiation. This switch may be coupled to a selection process, as the scFEA analysis revealed that many cells at 4 dpi could not sustain the metabolic changes triggered by cell activation. Finally, it occurs at a transition stage globally affecting the GRN, beyond metabolic regulation.

## 2. Methods

### 2.1 Data collection and processing

Sequencing reads were generated by Kurd et al. [33] and Milner et al. [34]. For these studies, the authors transferred P14, T-Cell-Ag-Receptor-transgenic CD8 T cells, which recognize a Lymphocytic ChorioMeningitis Virus epitope, to histocompatible hosts that were acutely immunized the day after with 10^5^ plaque-forming units of Lymphocytic ChorioMeningitis Virus Armstrong. Single responding P14 CD8 T cells from the spleens of immunized hosts were sorted at different days post infection (dpi) and loaded into Single Cell A chips for partition into Gel Bead In-Emulsions in a Chromium Controller (10x Genomics). Single-cell RNA libraries were prepared according to the 10x Genomics Chromium Single Cell 3′ Reagent Kits v2 User Guide and sequenced (paired-end) on a HiSeq 4000 (Illumina).

We used data collected at 0, 3, 4, 5, 6, 7, 10, 14, 21, 32 and 90 dpi and downloaded reads from GEO ‘release 2020-10-09’ (GSE131847) with fastqdump from the SRA-toolkit suite (https://www.ncbi.nlm.nih.gov/sra), using the --split-files option.

Quality control of reads was performed with fastp [35]: we discarded reads of too low quality, trimmed the others and removed adapters before selecting reads with a phred quality score [36] of at least 30.

Reads were then aligned to the mouse genome assembly GRCm38 (https://www.ncbi.nlm.nih.gov/assembly/GCF_000001635.20/) with the Kallisto-Bustools wrapper [37], using the ‘lamanno’ workflow to generate a table of Unique Molecular Identifier (UMI) counts of spliced mRNAs. Empty sequencing droplets were removed with a threshold set to the inflection knee of the curve representing the UMI counts of ranked droplets (http://bioconductor.org/books/3.15/OSCA.advanced/droplet-processing.html).

The UMI count tables of each collection day were imported and pooled in Seurat V4.0.4 [38] and the 14,666 genes that are expressed in at least 20 cells were selected. Dying cells with more than 5% of mitochondrial genes were further removed

The UMI count table was normalized with SCTransform [39] and 2% of the droplets containing more than one cell, as identified with the DoubletFinder package [40] were removed, resulting in an UMI count table of 14,666 genes expressed by 42,025 live single cells spanning the entire kinetics (Fig. 1A). Principal Component Analysis (PCA) was performed with prcomp (https://www.rdocumentation.org/packages/stats/versions/3.6.2/topics/prcomp). The broken-stick model [41] was used to select 9 principal components for further dimension reduction and visualization with Uniform Manifold Approximation and Projection (UMAP, Fig. 1B).

**Fig. 1.**
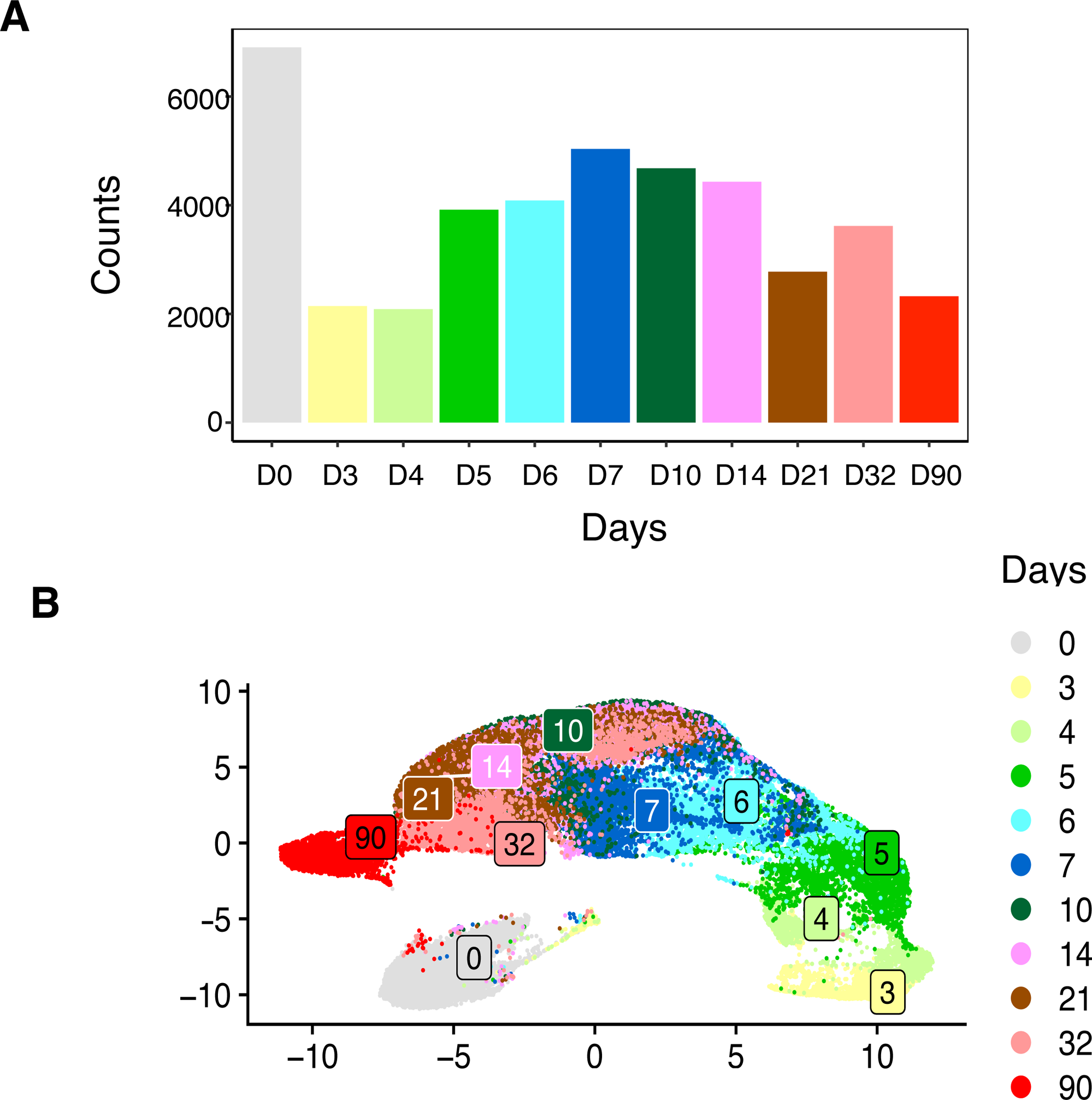
Data collection and processing. (A) Bar plot of live single cell counts recovered after selection at the indicated dpi. (B) UMAP representation of the recovered cells.

### 2.2 Metabolic gene selection

scFEA uses the expression of a list of 719 metabolic genes relevant to the estimation of the fluxes in all 171 metabolic modules. Among these metabolic genes, 430 are present in the 14,666 genes expressed in at least 20 cells in our data. However, all the genes regulating glycolysis and Oxphos described in Kyoto Encyclopedia of Genes and Genomes database (release ‘101.4’, Kanehisa and collaborators [71,72]) have not been included in the initial list. As metabolic switches from Oxphos to aerobic glycolysis and *vice versa* are crucial for the proper differentiation of CD8 T cells into effectors and memory cells during a viral infection, we complemented the list of 430 metabolic genes from N. Alghamdi et al. [30] with 4 and 96 extra genes, respectively involved in glycolysis and Oxphos, and present in the 14,666 considered genes (SupTable S1). We thus filtered the raw UMI count table for the expression of the 530 selected metabolic genes. Interestingly, the expression level of many metabolic genes displays temporal variations during the CD8 T cell response (SupFig. S1).

### 2.3 Data filtering and scFEA analysis

Raw UMI count data were filtered for metabolic genes in the designed list (SupTable S1) and the counts for 530 metabolic genes in the 42,025 cells were submitted to scFEA (version v1.1.2). The algorithm was installed and run at the High-Performance Computing cluster Pôle Scientifique de Modélisation Numérique of the Ecole Normale Supérieure de Lyon, according to the instructions of the authors (Alghamdi et al. [30], https://github.com/changwn/scFEA). It returned flux values in all cells for 168 modules.

### 2.4 Module selection for fluxome analyses

These 168 returned modules were filtered to select fluxes with a |CV|>7.10^−4^ for all cells to eliminate modules with too little flux variation along the kinetics (SupFig. S2). We checked the eliminated modules were not significantly varying during the CD8 T cell response and re-introduced the M-114 module that shows substantial variation with a |CV| of 2.10^−4^ (SupFig. S2D).

To only consider relevant flux values, the 131 resulting varying modules were then filtered to select those with a maximum flux value superior or equal to 10^−3^ AU. Among the 47 remaining modules, some still display weak flux values with a majority of cells not reaching the 10^−3^ AU threshold (SupFig. S3A). They correspond to modules with a median of values inferior to 3.10^−4^ AU (SupFig. S3B and C). These were filtered out and we finally removed from the 41 remaining modules, 7 modules corresponding to simple metabolite efflux from cells, resulting in a list of 34 modules of interest.

### 2.5 Metabolite selection

The concentration of the 70 metabolites, for which scFEA yielded results, were averaged by day and the delta between the highest and the lowest concentrations along the kinetics (concentration_delta) was calculated for each metabolite. The distribution of these concentration_deltas displays a gap around 10^−3^ AU (SupFig. S4) and we therefore selected the modules with a greater concentration_delta, ending up with 20 metabolites of interest. However, concentration in ornithine was hardly varying along the response (Fig. 5) and we did not consider it further, in the resulting top19 list (SupTable S2, SupFig. S5).

### 2.6 Gene expression variability

The variability of gene expression is estimated by the level of entropy of each gene at each dpi [44,45]. Briefly, for each dpi, the distribution of gene expression among the cell population is binned and Shannon’s entropy is then defined as minus the sum across bins of pk.log(pk), where pk is the probability for a cell to belong to bin k. Shannon’s Entropy thus measures heterogeneity in a population, with a value of 0 when all cells belong to the same bin (minimal entropy) and a maximal value of log(k)/k when they are evenly distributed in bins (maximal heterogeneity). The entropy of each metabolic gene at each dpi was estimated with the unbiased ‘best-upper-bound’ estimator, as in Paninski et al. [46].

### 2.7 Clustering of entropy kinetic profiles

To identify groups of genes with similar temporal entropy patterns, i.e. similar profiles of entropies along the kinetics, we used functional PCA followed by kmeans [47]. Briefly, functional PCA is used to reduce the dimensionality and catch temporal dynamics of entropy curves. We used a model with piece-wise constant functions (histograms) for their simplicity of interpretation. Then, a standard kmeans algorithm is used on principal components to find the appropriate clusters. The number of principal components and clusters (n = 2) was determined by the rule of thumb.

### 2.8 Gene Ontology analysis

The list of genes corresponding to the entropy kinetic profiles 1 and 2 in Fig. 6C, were submitted to DAVID Bioinformatic Resources (https://david.ncifcrf.gov/tools.jsp) for functional annotation clustering on Gene Ontologies ‘Molecular Function’, ‘Biological Process’ and ‘Cellular Component’. Seventy-eight out of the 86 genes in entropy kinetic profile1 supported the clustering in 10 annotation clusters with low stringency and 107 out of the 444 genes in entropy kinetic profile2 supported the clustering in 15 annotation clusters with high stringency (SupTable S3). The 78 and 107 genes supporting both annotation clusters were then used to map the corresponding enzymatic reactions on the KEGG pathway map (Fig. 7).

### 2.9 Cell clustering

The 2,089 cells collected at 4 dpi were selected and the UMI counts of the 14,666 genes were normalized with SCTransform [39]. The selected 2,100 highly variable genes were used for further dimension reduction with PCA and UMAP. Cell clustering was performed in Seurat [38] with a resolution level of 0.07 to obtain 2 clusters. Next, module flux values, as well as metabolite concentrations, in each D4-cell were normalized and clustered, with respective resolution values of 0.02 and 0.03 to again obtain 2 populations for each clustering.

### 2.10 Differential gene and gene set enrichment analyses

The 2,100 highly variable genes expressed by CD8 T cells at 4 dpi were selected and differential expression between cells from both clusters was evaluated by a kernel-based two-sample test that compares the distributions of gene expression [48]. P-values were adjusted by the Bonferroni method. The 111 and 47 genes up-regulated in gene expression clusters #0 and #1, respectively, with an average log2(FC) > 0.13 (SupTable S4) were then submitted to gene set enrichment analysis in g:Profiler (https://biit.cs.ut.ee/gprofiler/gost) with default values and the 2,100 highly variable genes used as the background list.

## 3. Results

### 3.1 Expression data

We obtained expression data from a scRNA-seq study of murine CD8 T cells responding to an acute viral infection [33]. Therein, the authors transferred P14 transgenic CD8 T cells that recognize a Lymphocytic ChorioMeningitis Virus epitope to histocompatible hosts, which were acutely immunized with the virus the day after. Single responding CD8 T cells from the spleens of immunized hosts were then sorted at different days post-infection (dpi) and analyzed by scRNA-seq. We performed quality control of the data downloaded from Gene Expression Omnibus (GSE131847) and selected cells and genes (2.1 Data collection and processing) to generate a final UMI count table of 14,666 genes in 42,025 single cells (Fig. 1A). Dimension reduction and representation on UMAP revealed a correct temporal arrangement of the cells (Fig. 1B), although D4 and D32 cells split in two regions of the projection.

### 3.2 Global analysis of metabolic changes in responding CD8 T cells

In order to have access to an unbiased global analysis of the metabolism in CD8 T cells responding to an *in vivo* viral infection at the single-cell level, we used the scFEA algorithm that can estimate metabolic fluxes from scRNA-seq expression data [30]. The authors have subdivided all the metabolic pathways of human and murine cells into 171 flux-independent modules of enzymatic reactions. These modules are then reconstructed as a factor graph based on the network topology and gene expression status. Finally, all cell fluxomes are estimated thanks to a multilayer neural network model to capture the nonlinear dependency of metabolic fluxes on the enzymatic gene expressions.

#### 3.2.1 Fluxome analysis

We selected 530 metabolic genes (2.2 Metabolic gene selection) for scFEA analysis. As expected, their level of expression varies during the primary T cell response (SupFig. S1). We submitted UMI counts for these 530 metabolic genes in the 42,025 cells to scFEA and the algorithm returned flux values in all cells for 168 of the designed modules (2.3 Data filtering and scFEA analysis). We selected modules with a substantial variation of flux along the differentiation kinetics and relevant flux values (2.4 Module selection for fluxome analyses), ending up with a list of 34 metabolic modules of interest (SupTable S5, SupFig. S6). Those belong to 13 of the 22 Super metabolic Modules (SM) defined by Alghamdi et al. [30], including SM1 corresponding to ‘glycolysis and TCA cycle’, which are crucial for CD8 T cell responses. Furthermore, spermine, beta-alanine, leucine+valine+isoleucine and spermine metabolisms; the pentose phosphate pathway; the urea cycle and the synthesis of purines, pyrimidines, N- and O-linked glycans, as well as of steroid hormones, all displayed significant and varying fluxes (SupTable S5).

Flux variations all occur between D0 and D10 of the response with a surge in variation at D5, all fluxes returning to their initial values by D10 and remaining stable up to the memory phase of the response at D90 (SupFig. S6). The flux variations display 3 typical patterns: fluxes either increase (Fig. 2A) or decrease (Fig. 2B) between D0 and D10, or show a biphasic evolution with a transient decrease between D0 and D4, followed by an increase above initial values from 5 to 10 dpi (Fig. 2C). Whatever the direction of the flux variations, cells at 4 dpi are heterogeneous with some of the cells displaying a flux corresponding to the variation between 3 and 5 dpi, but others with a flux similar to initial values, as if they were not able to sustain the initial metabolic changes.

**Fig. 2.**
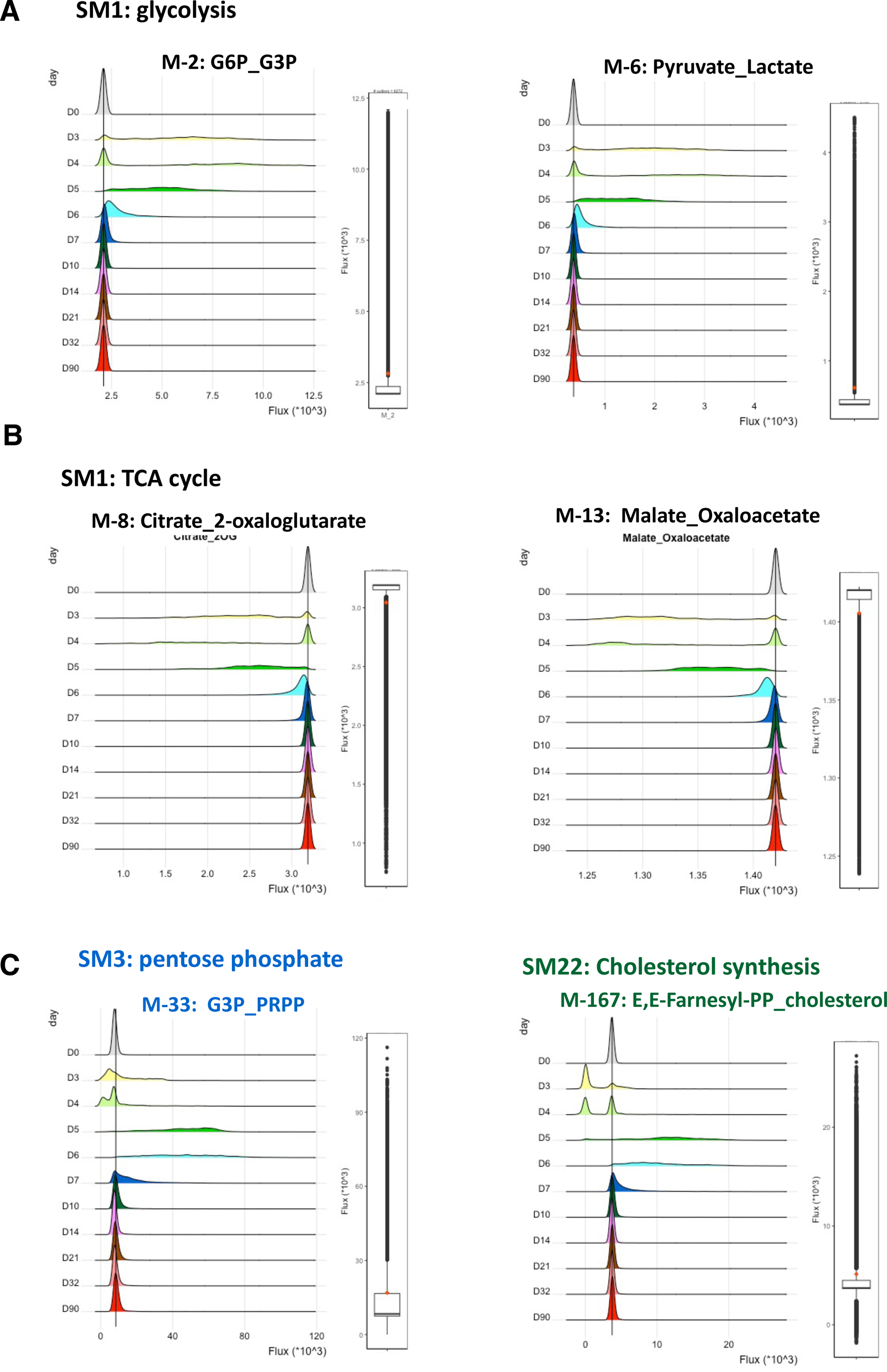
Examples of module flux variations. For each metabolic module, the distributions of flux values (x10^3^ AU) at all dpi are shown left-hand and a box-plot of flux values for all cells is shown right-hand. The vertical bar on the left and the red dot on the right indicate the median and mean of flux values, respectively. Examples of modules with an upregulated (A), downregulated (B) or biphasic (C) flux are shown. Names, as well as the initial and final compounds, of modules are indicated in the color of the corresponding Super-Module, as in Fig. 3.

The projection of the fluctuating flux modules on the murine metabolic map from KEGG reveals a regionalization of the selected modules by Super-Module (Fig. 3) and complex variations of fluxes during the differentiation of cells. For instance, in the ‘Pyrimidine synthesis’ SM21, the flux through M-155 that corresponds to the reactions Uridine-TriPhospsate (UTP) → Cytidine-TriPhosphate (CTP) → Cytidine-DiPhosphate (CDP) is upregulated between D0 and D10, while the flux through M-153 that corresponds to the overlapping reaction Uridine-MonoPhosphate (UMP) → UTP → CTP → CDP is initially downregulated between D0 and D4 and then upregulated (Fig. 4A). Furthermore, the fluxes through M-157 (CDP → deoxyCDP (dCDP)), M-158 (dCDP → deoxycytidine) and M-171 (dCDP → deoxycytidine-TriPhosphate (dCTP)) are up regulated, while the flux through M-161 (dCDP → deoxycytidine-MonoPhosphate (dCMP) is downregulated. Such complex flux interactions are difficult to interpret in terms of metabolite concentrations, such as for CDP or dCDP, and were evidenced in other Super-Modules, such as SM17 and SM16 (Fig. 4B).

**Fig. 3.**
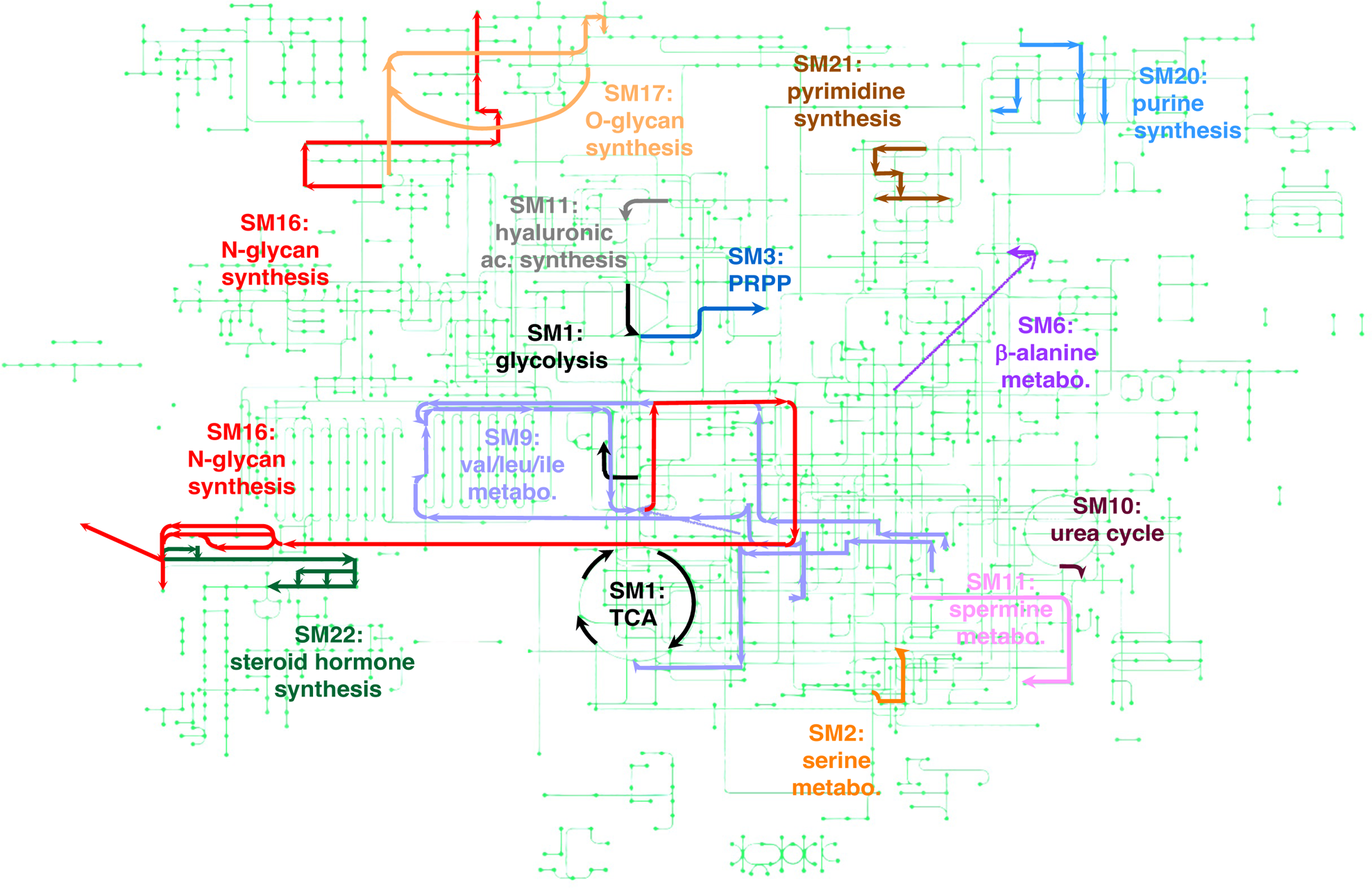
Metabolic map projection of the modules selected for analysis. The modules are depicted on the global murine metabolic pathway map ‘mmu01100’ from KEGG [42,43]. They are represented as arrows following the enzymatic reactions from their initial to the terminal compounds. Modules are colored according to their SM family as in [30]. Detailed analysis of module fluxes and metabolite concentrations are depicted in Fig. 4C-H.

**Fig. 4.**
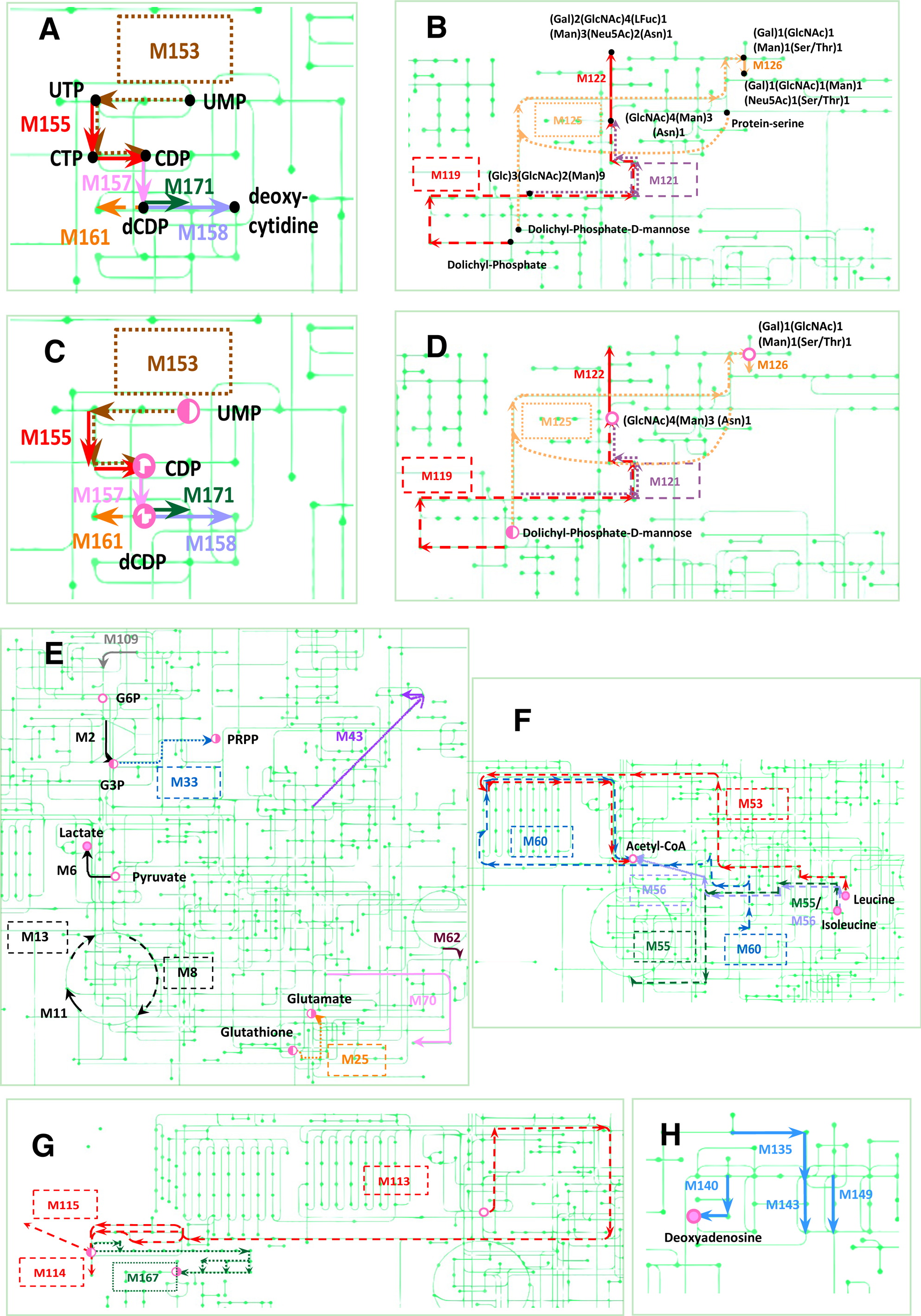
Detailed metabolic map projection of the modules and metabolites selected for analysis. The modules are depicted on parts of the global murine metabolic pathway map ‘mmu01100’ from KEGG [42,43]. They are represented as arrows following the enzymatic reactions from their initial to the terminal compounds. Plain arrows and unboxed names correspond to fluxes upregulated between D0 and D10. Dashed arrows and name boxes correspond to fluxes downregulated between D0 and D10. Dotted arrows and name boxes correspond to fluxes transiently downregulated between D0 and D4 and upregulated between D5 and D10. (A-B) Intermediate compounds are indicated by black dots and labels. (A) For the sake of clarity, the selected modules of SM21 (pyrimidine metabolism) are represented in a different color: the flux through M155, M157, M158, M161 and M171 is up-regulated, while down-regulated in M153. (B) Selected modules of SM16 (N-linked) and SM17 (O-linked) glycan synthesis super-modules are presented in different colors. In SM16, the flux through M119 is down-regulated, up-regulated through M122 and fluctuates for M121. In SM17, flux is fluctuating for M125 and up-regulated in M126. (C-H) Only the top19 variable metabolites are labelled and shown as pink circles. Empty/plain circles represent respectively metabolites with a transient depletion/accumulation between D0 and D14. Metabolites with a biphasic change in concentrations are represented with two-color circles, with a pink left-hand part for metabolites firstly accumulated and with a pink right-hand part for those that are first depleted from cells. (C) As in panel A, with selected metabolites. CDP and dCDP with a triphasic variation are represented by a hatched circle. (D) As in panel B, with selected metabolites. (E) Glycolysis and TCA-cycle SM1 is represented in black, with the flux up-regulated in M2, M6 and M11 and down-regulated in M8 and M13. Unique modules were selected from SM2 (serine metabolism), SM3 (PPP), SM6 (beta-alanine metabolism), SM10 (Urea cycle), SM11 (spermine metabolism) and SM13 (Hyaluronic acid synthesis) super-modules. They are represented in the corresponding color, as in Fig 3. The first reactions of M43 are not depicted in the metabolic map and are represented as a direct jump. (F) For the sake of clarity, the selected modules of SM9 (leucine, valine and isoleucine metabolism) are represented in different colors. The flux through all selected modules of SM9 is down-regulated. The last reactions to acetyl-coA for module M-56 are represented as a direct jump the intermediate chemical reactions are not depicted on the metabolic map. (G) Selected modules of SM16 (N-linked glycan synthesis) and SM22 (steroid hormone synthesis) are represented in their respective color, as in Fig. 3. The flux through all modules of SM16 is down-regulated. Note that module M-115 is not pointing to its final product Farnesal, which is not represented on the map. The flux is fluctuating through M167 of SM22. (H). The flux through all selected modules of the SM20 (purine synthesis) super-module is up-regulated.

#### 3.2.2 Metabolic stress analysis

Thus, in order to relate flux variations in modules and metabolite accumulation or deprivation from cells, we again used scFEA that calculates metabolite concentrations from the estimated fluxes. scFEA returned values for 70 metabolites. We selected (2.5 Metabolite selection) the top19 variable metabolites for analysis (SupTable S2, SupFig S5). The concentration of metabolites varied between D0 and D14 of the response (Fig. 5) with a peak at 5 dpi. Cells were transiently deprived of Glucose-6-Phosphate (G6P), pyruvate, acetyl-coA and glycan backbones (GlcNAc)4 (Man)3 (Asn)1 and (Gal)1 (GlcNAc)1 (Man)1 (Ser/Thr)1; and transiently accumulated leucine, isoleucine, lactate and deoxyadenosine. For these metabolites, a large fraction of cells at 4 dpi did not sustain the accumulation or loss, as observed for fluxes (Fig. 2). Other metabolite concentrations were biphasic with G3P, glutathione, E,E-Farnesyl-PP, UMP and dolichyl-P-D-mannose being accumulated up to 4 dpi and then deprived from cells. Conversely, after an initial loss from D0 to D4, cells accumulated glutamate, cholesterol and 5-Phosphoribose-2P (PRPP) between D5 and D14. Finally, CDP concentration displayed oscillations rising at D3, then decreasing below initial values with a nadir at 5 dpi, followed by a new maximum at D6, while dCDP concentration oscillated in mirror (Fig. 5).

**Fig. 5.**
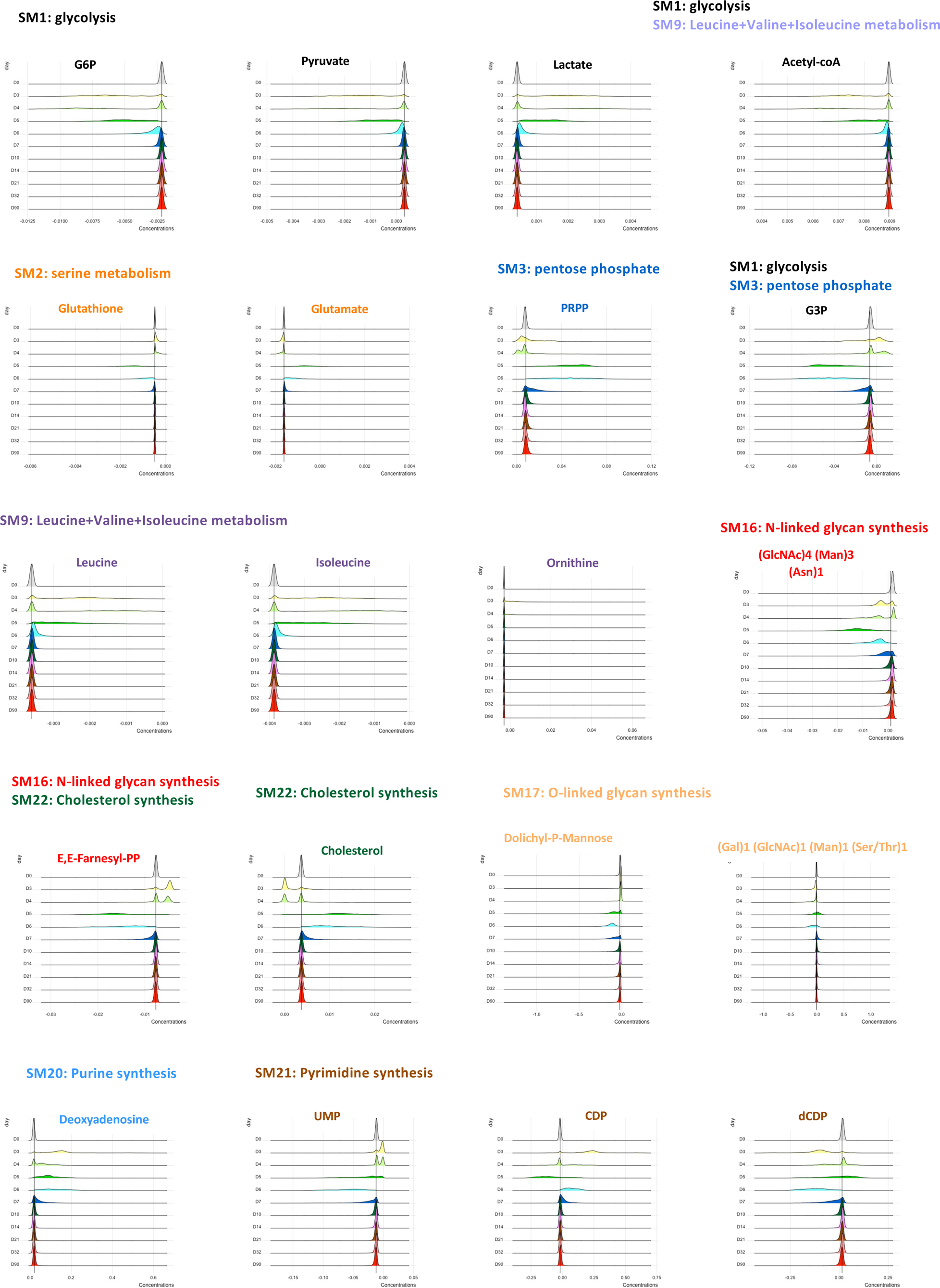
Variation in concentration of the top20 most variables metabolites. For each metabolite, the distributions of concentrations (AU) at all dpi are shown. The vertical bar indicates the median for all cells. Corresponding Super-Modules are indicated in their respective color. For metabolites at the junction of 2 SM, both are indicated.

Interestingly, when the top19 variable metabolites were localized on the metabolic map, they all matched a varying flux module (Fig. 4). Indeed, Glutamate, PRPP and cholesterol, which are the products of M-25, M-33, M-167 modules with first decreasing and then up-regulated fluxes, were first deleted from cells before accumulating, while Glutathione, G3P and (E,E)-Farnesyl-PP, the substrates of these same modules were first accumulated before loss (Fig. 4E). Also, leucine and isoleucine substrates of M-53 and M-55/M-56 modules with decreasing fluxes (Fig. 4F) and deoxyadenosine product of M-140 (Fig. 4H) with an increasing flux accumulated, while the increase of M-6 flux from pyruvate to lactate resulted in an accumulation of lactate and pyruvate depletion (Fig. 4E). Furthermore, the analysis of metabolite stress helped interpret the complex interactions of varying fluxes. For instance, the complex interactions of fluxes in all 6 modules of the ‘pyrimidine synthesis’ SM21 resulted in opposite oscillations in CDP and dCDP concentrations, while fluctuations in M-153 flux resulted in an accumulation of UMP prior to depletion (Fig. 4C). Similarly, the rise in M-126 flux counteracted fluctuations in M-125 flux and the increase in M-122 competed the decrease and fluctuations of M-119 and M-121, so that cells were depleted from (Gal)1 (GlcNAc)1 (Man)1 (Ser/Thr)1 and (GlcNAc)4 (Man)3 (Asn)1 glycans (Fig. 4D). This is in accordance with the stronger values of fluxes in M-126 and M-122, as compared to fluxes in M-125 and M-119 and M-121, respectively (SupTable S5).

In a whole, scFEA analysis led to a very detailed global and temporal single-cell description of variations in metabolic fluxes and metabolite concentrations in CD8 T cells during a viral infection.

### 3.3 Cell-to-cell variability in metabolic gene expression

Single-cell resolution of metabolic fluxome analysis showed a large degree of inter-cellular heterogeneity, revealing a cell subset at 4 dpi (Fig. 2, SupFig. S6) and highlighting the importance of this degree of granularity to characterize CD8 T cell responses. We next questioned whether inter-cellular heterogeneity in metabolic fluxes stems from a certain degree of stochasticity in the differentiation process. Indeed, in most cell differentiation contexts, a rise in the stochasticity of gene expression has been observed just prior to transition or branching points in the differentiation process [49,50,45,51,52,31]. Such an increase in gene expression variability results in increased inter-cellular variability and allows cells submitted to environmental perturbations to explore the gene expression space before selection of the most appropriate GRN state in the resulting differentiated population [53,54]. This phenomenon has not yet been described for mature lymphocytes differentiating in response to an antigenic challenge. As shown in SupFig. S1, the level of metabolic gene expression varies during CD8 T cell differentiation. However, during hematopoiesis, many genes show fluctuations in expression level, but pathway-specific genes are the ones that show the highest fluctuations in cell-to-cell variability over the course of a differentiation trajectory [45]. Since metabolic regulation is crucial for the proper differentiation of CD8 T cells in response to a viral challenge, we thus assessed the inter-cellular variability in metabolic gene expression during this differentiation process.

For this, we estimated the degree of inter-cellular variability by the level of entropy [44,45] for the 530 metabolic genes selected for flux estimations, at each dpi (2.6 Gene expression variability). As shown in Fig. 6A, many metabolic genes show strong variations in expression variability, mostly during the first 10 days after infection. Thus, although metabolic gene expression is strictly regulated during CD8 T cell differentiation [5,24], it remains submitted to surges in inter-cellular variability, which suggests the process is submitted to variation in stochasticity. There is a weak correlation (r=0.68, Pearson’s coefficient) between the variations in gene expression and entropy (SupFig. S7), as previously seen in other cell differentiation models [45]. In order to highlight kinetic patterns of metabolic gene entropies, we clustered entropy profiles shown in Fig. 6A, with functional PCA followed by kmeans clustering (2.7 Clustering of entropy profiles). Thus, we could separate kinetic patterns of entropies into two groups (Fig. 6B and C): 86 genes (SupTable S6) show a strong, transient and immediate surge in entropy after activation, hereafter called profile 1, while 444 genes show a weaker and later increase in entropy, beginning at 4 dpi, with a peak at 6 dpi, hereafter called profile 2.

**Fig. 6.**
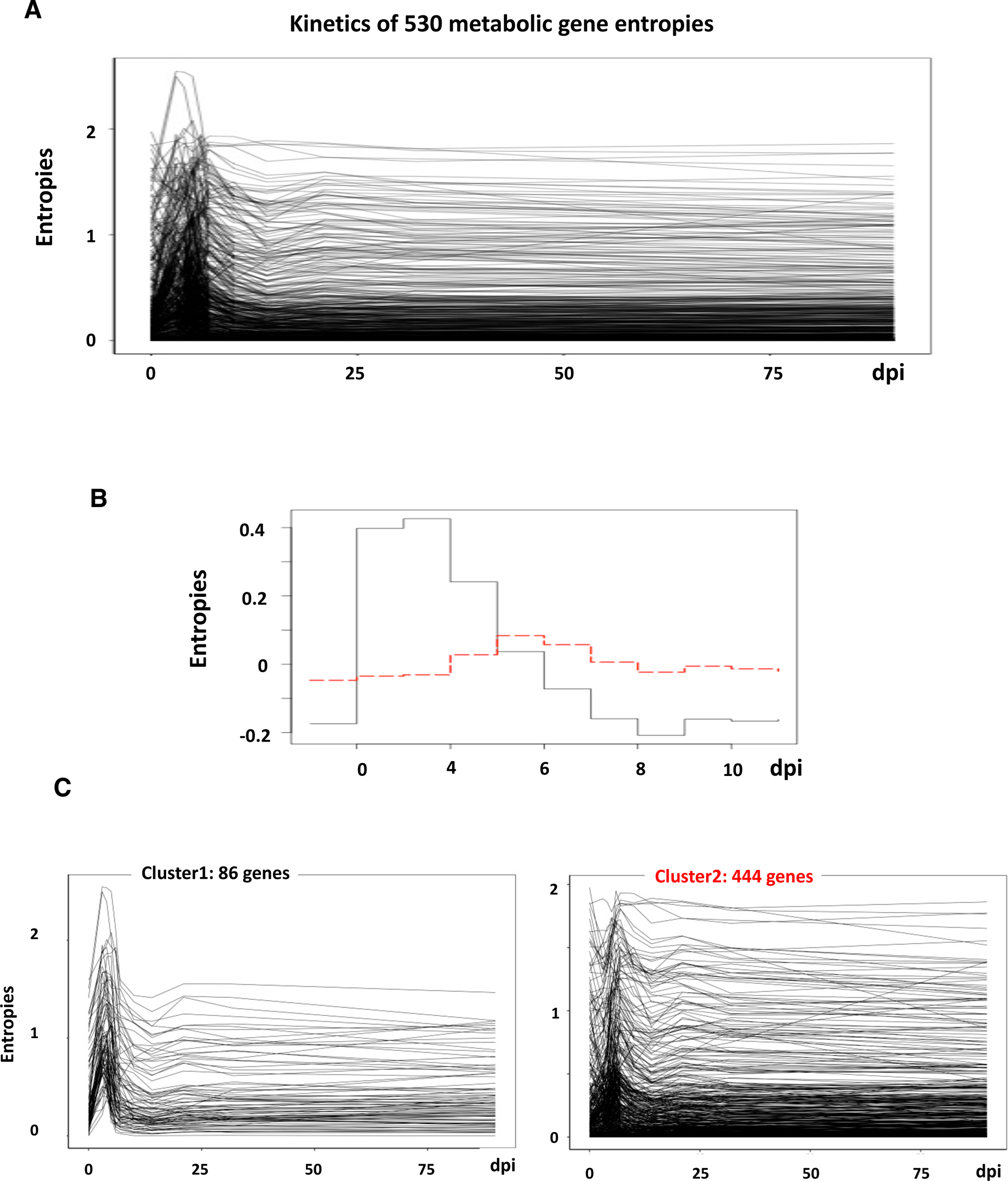
Metabolic genes expression inter-cellular variability. (A) The entropies of the 530 metabolic genes were calculated at each collection day and are represented as a function of time. Note that we draw lines between time points for visualization purposes only, as we cannot assume monotonous variation between 2 distant measures, such as D32 and D90. (B) Functional PCA followed by kmeans clustering evidenced 2 groups of kinetic patterns. (C) The kinetics of the entropies of the genes in both clusters of panel B are shown as in panel A.

We next investigated the metabolic pathways sustained by the genes submitted to variation in inter-cellular variability at these two stages. For this, we performed a Gene Ontology analysis of both groups of genes (2.8 Gene ontology analysis) and mapped the reactions they govern on the KEGG global metabolic map. As shown in Fig. 7, the metabolic pathways corresponding to the genes allowing the functional annotation clustering of entropy kinetic profiles 1 and 2 cover a great deal of the murine metabolic map and hardly overlap (See also ‘GO terms’ in SupTable S3). Different sections of the CD8 T cell metabolism are thus affected by the early and late increases in gene expression variability.

**Fig. 7.**
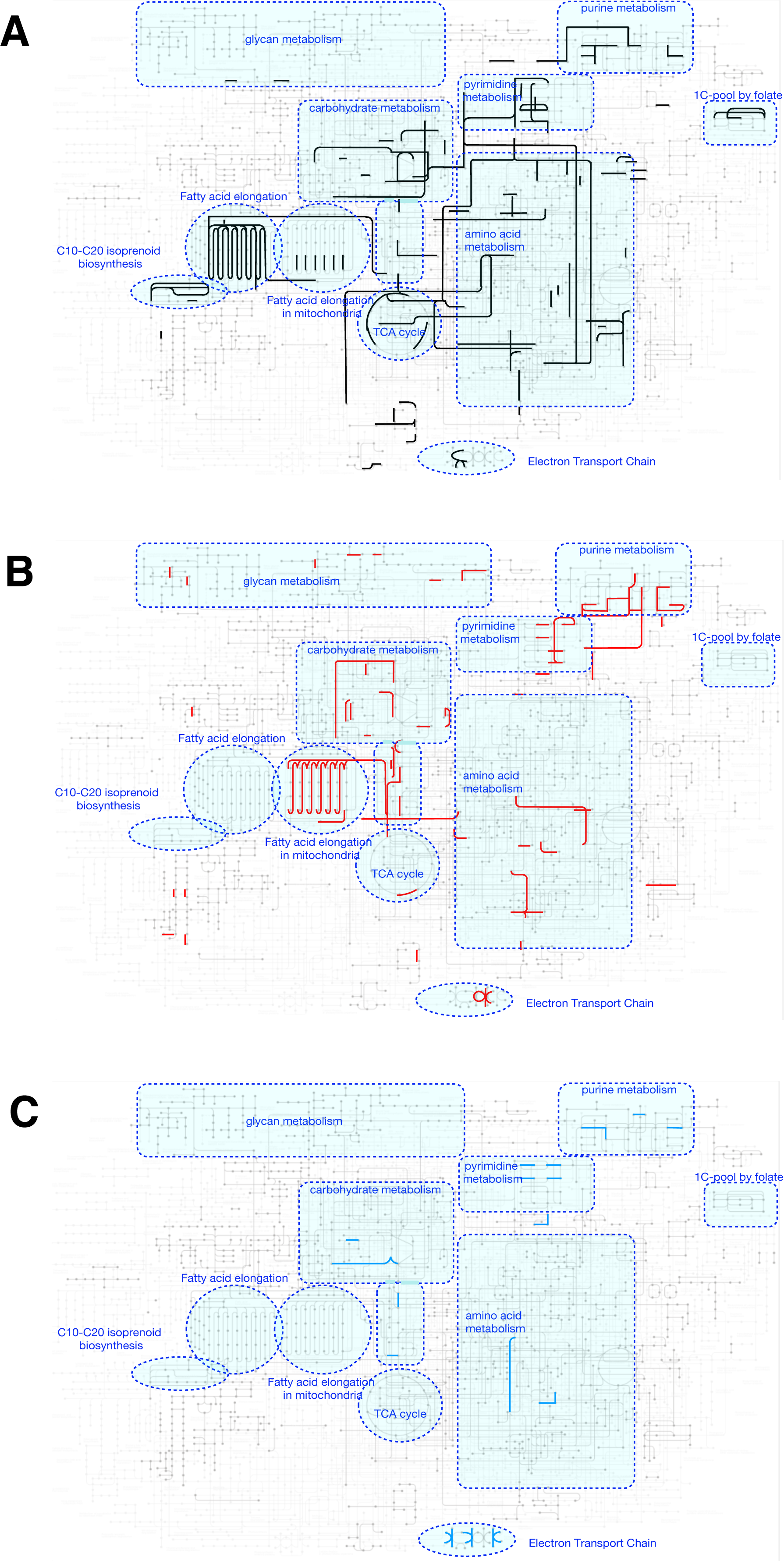
Metabolic map projection of the pathways covered by genes in kinetic profiles. The enzymatic reactions corresponding to genes used for functional annotation of kinetic profiles are depicted on the KEGG murine metabolic map in (A) black for immediate profile 1, (B) red for later profile 2 and (C) blue for both. The regions corresponding to large pans of cellular metabolism are highlighted and labeled in blue.

### 3.4 Metabolism analysis reveals a transition stage 4 days post-infection

The late surge in gene expression inter-cellular variability suggests a new transition step at 4 dpi that may be responsible for the heterogeneity in fluxes and metabolite concentrations observed at that time of the response (Fig. 2 and Fig. 5). Indeed, we clustered cells harvested at D4 based on the expression of 2,100 highly variable genes (Fig. 8B) and it is striking that the dichotomy observed at 4 dpi in module flux values and metabolite concentrations largely overlaps the clustering (2.9 Cell clustering) obtained based on all highly variable gene expression (Fig. 8A and data not shown). Furthermore, when cells collected at 4 dpi are clustered based on flux module values or metabolite concentrations, a large overlap is again observed between these clusters and those based on all highly variable gene expression data (Fig. 8 B and C). Thus, the late surge in inter-cellular metabolic gene expression variability (Fig. 6 B and C) reveals a transition stage affecting the whole GRN state and specifically metabolic activity, and points toward the existence of two distinct cell populations at that stage. Gene set enrichment analysis (2.10 Differential gene and gene set enrichment analyses) of D4-expression clusters mostly revealed an enrichment in genes associated with CD8 T cell effector functions (e.g., Ccl3, Ccl4, Ccl5, Ifng, Gzmb, Prf1) in cells from expression cluster #0 and the specific enrichment in “Zf5” motif of the Zbtb4 transcription repressor in 32 out of 47 over-expressed genes in cells from expression cluster #1 (SupTable S7).

**Fig. 8.**
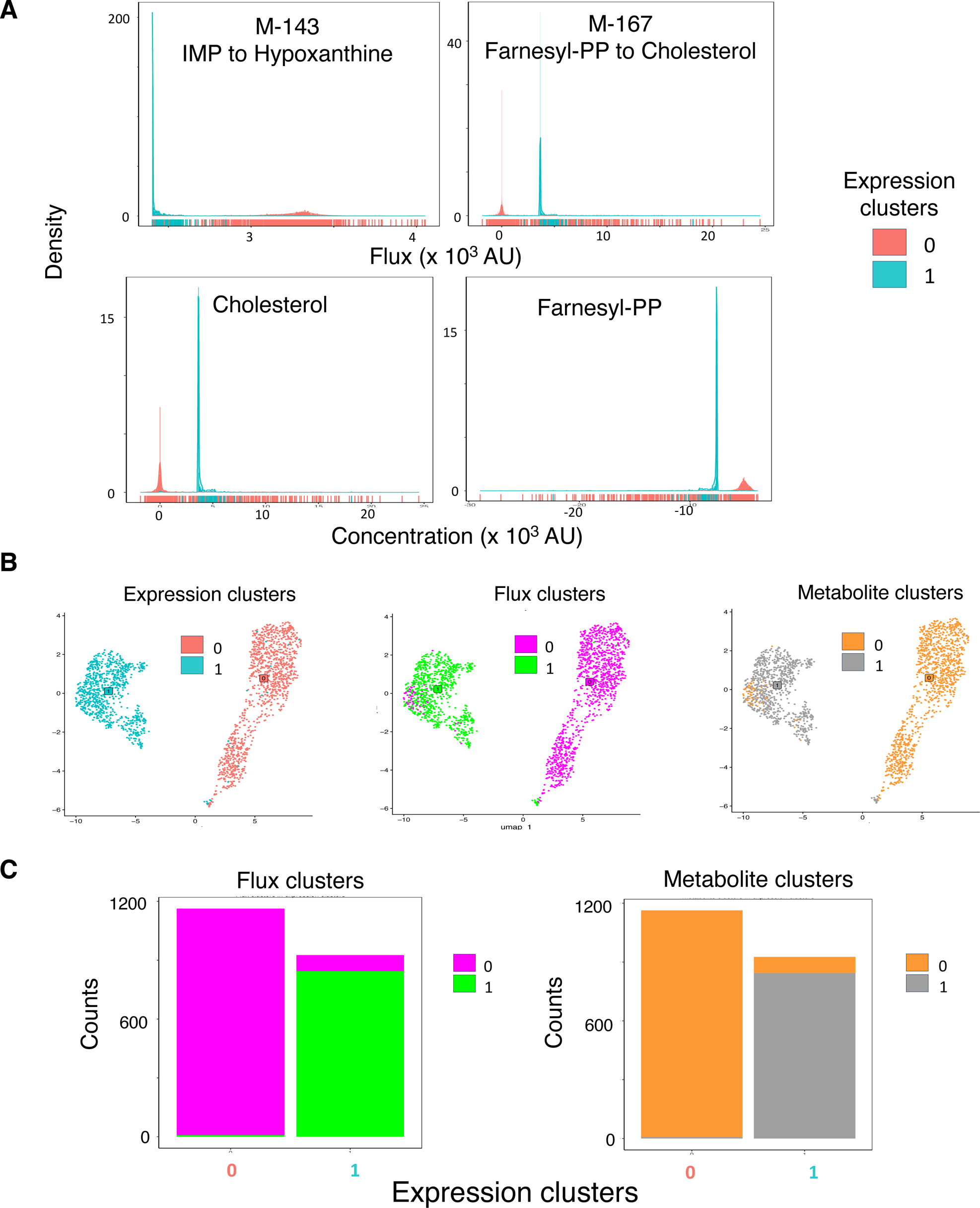
Module flux values and metabolite concentration of clustered D4 cells. (A) Cells collected at 4 dpi were clustered on their expression levels of 2,100 highly variable genes and the histograms of flux values of two modules (top) and two metabolite concentrations (bottom) are colored according to the expression clusters #0 (red) and #1 (blue). (B) UMAP of D4 cell expression data colored by expression clusters (left, #0 in red and #1 in blue), flux module value clusters (middle, #0 in pink and #1 in green) and metabolite concentration clusters (right, #0 in orange and #1 in grey). (C) Cell repartition of flux module value clusters (left, #0 in pink and #1 in green) and metabolite concentration clusters (right, #0 in orange and #1 in grey) within expression data clusters.

## 4. Discussion

In this study we have made use of the scFEA algorithm [30] to evaluate at the single-cell level the global metabolic activity in CD8 T cells responding to a viral infection *in vivo* from scRNA-seq data [33,34]. Previous single-CD8-T-cell metabolic analyses relied only on scRNA-seq measurements of mRNA levels of *in vitro* activated cells [27], or cytometry quantification of a few selected proteins implied in metabolism and/or its regulation [25,55,56]. However, mRNA and protein levels are not linearly correlated to the activity of the corresponding metabolic modules and our study is the first attempt at globally evaluating all metabolic activities in single responding CD8 T cells. The conclusions of our study can be questioned as they are drawn from flux value estimations from expression data. Two lines of evidence nevertheless support our claims: first, the output of the scFEA algorithm was validated against experimental data in the original study [30] and second, our analysis does reproduce known facts on metabolic changes in activated CD8 T cells, demonstrating the validity of the approach. Indeed, flux estimation of carbohydrate metabolism pathways was in accordance with the extensively described switches in glucose consumption observed during CD8 T cell differentiation [7,3–5]. Among the five modules describing the degradation of glucose to pyruvate, only M-2 (Glucose-6P (G6P) to G3P) showed a substantial flux variation but its flux increase upon activation is in agreement with a global rise in metabolic activity of antigen-challenged CD8 T cells. Furthermore, the flux increase in M-6 (pyruvate to lactate) exactly describes the metabolic switch from Oxphos to aerobic glycolysis that is crucial for CD8 T cell differentiation into effector cells. Oxphos is not entirely included in modules of the scFEA analysis, however the flux through M-8 (citrate to 2-oxoglutarate) and M-13 (malate to oxaloacetate) of the TCA, which fuels Oxphos, are decreased as expected. Of note, varying fluxes through modules may seem contradictory and sometimes difficult to interpret. However, metabolic stress analysis helped reconciling the observations on fluxomes. For instance, if the conflicting increase in M-11 (succinate to fumarate) is difficult to interpret directly in regards to M-8 and M-13 modules, metabolic stress analysis revealed a depletion of acetyl-coA that, together with the deprivation of G6P and pyruvate, and with the accumulation of lactate, all confirm the metabolic switch from Oxphos to aerobic glycolysis during the differentiation of virus-challenged CD8 T cells into effectors. Besides, the increased flux through M-62 (ornithine to putrescine) corresponds to the increase in polyamine synthesis from ornithine through the urea cycle, as described in Ag-activated T cells [18]. scFEA analysis also added some temporal information on other known CD8 T cell metabolic perturbations. For instance, glucose consumption through the pentose phosphate anabolic pathway is increased in activated CD8 T cells [18,57]. It is thought to be important for nucleic acid demands in highly proliferating cells [7,58]. We observed that the flux through M-33 (G3P to PRPP) of the PPP initially decreases leading, together with the increase flux through M-2, to an accumulation of G3P at 4 dpi. The increase in M-33 flux leading to an accumulation of PRPP at D6, in only seen secondary to the initial decrease, suggesting an important role for the PPP during the transition from activated cells to fully differentiated effectors. Similarly, T cell activation induces the increase of glutamine uptake and glutaminolysis [17,18]. The glutamate produced by glutamine degradation can enter mitochondria and fuel the TCA to produce α-KG and downstream cellular building blocks, such as lipids and nucleic acids [4,10,59,60], or be converted to glutathione in the cytoplasm to reduce the Reactive Oxygen Species produced by the enhanced mitochondrial activity [3,5]. We observed an initial decrease in the flux through the M-25 (glutathione to glutamate), followed by an increase which led to accumulation of glutathione at D3 and glutamate at D5. This suggests that increased glutaminolysis is initially used to produce glutathione that will buffer the Reactive Oxygen Species production by mitochondria, in turn sustaining the mammalian Target Of rapamycin pathway and the glycolytic switch [61]. After 4 dpi, enhanced glutaminolysis leads to glutamate accumulation that can fuel the TCA cycle to produce *de novo* cellular components in highly proliferating effector cells [4,10,59,60]. Finally, we detected the increased cholesterol production described in activated CD8 T cells [15,16] only after 4 dpi, suggesting its role may not be major for the immune synapse formation [62] but important for the regulation of the transcriptional activity of Liver X receptor [63] and Sterol Regulatory Element-Binding Protein [16], in activated T cells [5]. In conclusion, scFEA analysis provided a very detailed description of metabolic adaptations in virus activated CD8 T cells at the single cell level that were validated by previous experimental results but added a level of temporal precision.

Nevertheless, no substantial variations of fluxes or metabolite concentrations in cells were observed past 10 dpi and, thus, the back-switch in metabolism from aerobic glycolysis to Oxphos and FAO of effectors differentiating into memory CD8 T cells [3,24,5] was not evidenced by our analysis. This may be due to variations in flux and metabolite concentrations that are beyond the sensitivity of scFEA estimations. scFEA could not either detect flux variations in any of the modules of the SM4, 5, 7, 8, 14, 15, 18 or 19, defined in Alghamdi et al. [30] as ‘glycan, glycosaminoglycan, glutamate, glycogen, sialic and fatty acids, beta-alanine and aspartate metabolism or syntheses’. We cannot conclude whether these sections of the CD8 T cell metabolism are less affected by cellular differentiation or whether the algorithm fails to evidence them. However, as many genes encoding enzymes responsible for reactions of some of these pathways, such fatty acid elongation, are submitted to a surge in expression variability, we suspect scFEA is not sensitive enough to catch all CD8 T cell metabolic perturbations, further strengthening the quantitative importance of the changes that were detected. In that respect, scFEA revealed variations of full sections of cellular metabolism that were not expected from experimental studies. For instance, scFEA revealed perturbations of the fluxomes in ‘N-and O-linked glycan synthesis’ SM16 and SM17 with deprivation of two types glycan backbones. Interestingly, enhanced glycolysis and glutaminolysis fuel an increase in O-linked GlcNAcylation of nuclear proteins upon T cell activation [64,65] that triggers the Nuclear Factor kappa-light-chain enhancer of activated B cells signaling pathway [66]. The differential regulation of specific modules and metabolites in SM16 and SM17 highlighted by scFEA could help designing experiments to decipher which post-translational modifications are crucial for CD8 T cell antiviral responses. Furthermore, reduced flux in several modules of leucine and isoleucine degradation to succinyl-coA and acetyl-coA in SM9 resulted in the accumulation of both these amino acids and participated with aerobic glycolysis to the deprivation in acetyl-coA. Thus, scFEA suggests a role for these amino acids in CD8 T cell responses that remains to be experimentally investigated. Deprivation of acetyl-coA, the principal giver of acyl groups for post-translational modifications [67] shall also deserve experimental investigation. Similarly, scFEA analysis revealed an increase in the flux through several modules of the ‘purine synthesis’ SM20, leading to the accumulation of deoxyadenosine, as well as a very complex interaction of fluxomes in modules and variations of metabolite concentrations in the ‘pyrimidine synthesis’ SM21. The role of specific intermediates and metabolic modules of nucleic acid biosynthesis in CD8 T cell activation is totally unexplored. In conclusion, scFEA analysis points out new sections of cellular metabolism that deserve experimental investigation in the context of CD8 T cell antiviral responses and memory development. Our observations were based on the *in vivo* response to the highly studied LCMV virus. Those that were described for bulk populations in many experimental models [3–5, 7–9] are probably common to all acute anti-viral responses. It would be very interesting that experimental validation of detailed temporal and new observations addresses, which are common or pathogen-specific during CD8 T cell responses.

Cell differentiation has long been seen as an instructive process with genetic programs orchestrated by a set of master regulator genes, the transcription factors, being solicited in every single cell submitted to environmental perturbation [68–70]. The recent breakthrough of single cell omics-technologies has largely challenged this view [71]. Indeed, stochasticity of gene expression at the single cell level [72] translates to stochasticity in the differentiation process [73,74]. In this Darwinian view, cell differentiation proceeds in two steps: stochastic gene expression in response to a stimulus creates a certain degree of transcriptional uncertainty, where individual cells can initiate different genetic programs, leading to a high degree of inter-cell variability in gene expression. The subsequent selection of fit cells allows the return to a homogeneous population of differentiated cells, in which a new stable state of the GRN has been established [75,49,50,45,51,52,31].

Although this differentiation mechanism has been demonstrated in all biological models examined to date [45,52], we show here for the first time that metabolic gene expression during an *in vivo* lymphocyte response to a viral challenge is submitted to surges in inter-cellular variability. Inter-cellular gene expression level variability does not uniquely result from stochastic gene expression, but genes involved in differentiation commitments show a high degree of inter-cellular variability [45]. Thus, our results show that, as many other differentiation processes, CD8 T cell responses might benefit from a stochastic exploratory search of the gene expression space during their differentiation at least at the level of the metabolic functions.

This implies the existence of selection steps of cells with the best fit state of the GRN at some points of the process. Our observations suggest that such a selection step may occur around 4 dpi. Indeed, at this stage a large fraction of cells seems to recover flux values and metabolite concentrations similar to those in unstimulated cells, as if unable to sustain the metabolic changes occurring in other cells between D3 and D5 of the response. Furthermore, in many cases the flux direction in modules, as well as the corresponding metabolite accumulation/deprivation in cells, reverts between D3 and D5 highlighting a metabolic switch at D4. In addition, kinetic patterns of gene expression variability revealed two groups of gene controlling different sections of the cellular metabolism, that show a surge in entropy either directly after infection, or later starting at 4 dpi, suggesting a transition stage at that time of the response. This is strengthened by the observation that heterogeneity of metabolic activities in cells collected at 4 dpi matches different states of the global GRN. Altogether, these results suggest a transition stage accompanied by a metabolic switch in activated cells around 4 dpi, when they start to differentiate into cytotoxic effectors, that would result in the selection of the best metabolically fit cells.

## 5. Conclusions

In a whole, our study allowed a global survey of cellular metabolism in CD8 T cell responding to a viral infection *in vivo*, at the single-cell level, highlighting metabolic perturbations beyond known switches revealed by previous experimental approaches. This study also evidenced a transition step around 4 dpi, accompanied by a metabolic switch during the differentiation of antigen-activated cells into full-blown effectors, adding one more piece of evidence linking metabolic activity variations and differentiation processes.

## Acknowledgments

Christophe Arpin warmly thanks Matteo Bouvier and Maxime Lepetit for their help in efficient coding. The authors wish to thank Drs. John T Chang [33] and Chi Zhang [30] for providing helpful information on their data and algorithm, respectively. We gratefully acknowledge support from the Pôle Scientifique de Modélisation Numérique of the Ecole Normale Supérieure de Lyon, especially Loïs Taulelle, and the Institut Français de Bioinformatique for the computing resources. We also thank the bioinformatic hub of the Laboratory of Biology and Modeling of the Cell (LBMC), especially Laurent Modolo, for their teaching and help. We thank the BioSyL Federation and the LabEx Ecofect (ANR-11-LABX-0048) of the University of Lyon for inspiring scientific events. We warmly thank Dr. Jacqueline Marvel for critical reading of the manuscript.

## CRediT authorship contribution statement

**Christophe Arpin:** Conceptualization, Methodology, Validation, Formal Analysis, Software, Investigation, Data Curation, Writing: Review and editing, Writing: Original draft preparation and Project Administration. **Franck Picard:** Formal Analysis, Writing: Review and editing. **Olivier Gandrillon:** Conceptualization, Methodology, Validation, Writing: Review and editing, Supervision Project Administration and Funding Acquisition.

## Funding

This research was supported by a recurrent funding from Centre National de la Recherche Scientifique (CNRS), France, and did not receive any specific grant from funding agencies in the public, commercial, or not-for-profit sectors.

## Abbreviations

FAO: FA oxidation
G3P: Glyceraldehyde-3-Phosphate
G6P: Glucose-6-Phosphate
GRN: Gene Regulatory Network
KEGG: Kyoto Encyclopedia of Genes and G
PPP: Pentose Phosphate Pathway
PRPP: 5-Phosphoribose-2P
scFEA: single-cell Flux Estimation Analysis
scRNA-seq: single-cell RNA-sequencing
SM: Super metabolic Module
TCA: TriCarboxylic Acid
UMI: Unique Molecular Identifier

## Notes

### Competing Interest Statement

The authors have declared no competing interest.

### Summary of Updates

Updated after review process. A new Supplementary figure (SupFig. S5) has been added and previous over loaded fig.3 has been split in Fig. 3 and Fig.4

