## Supplementary material for "*In silico* single-cell metabolism analysis unravels a new transition stage of CD8 T cells 4 days post-infection"

### Supplementary information

#### Supporting Figures

- SupFig. S1 Metabolic gene longitudinal expression.
- SupFig. S2 Selection of flux modules with substantial variation.
- SupFig. S3 Selection of modules with relevant flux values.
- SupFig. S4 Distribution of metabolite concentration_deltas.
- SupFig. S5 Variations of the concentrations of the top19 variable metabolites.
- SupFig. S6 Fluxome representation of all the 34 modules selected for analysis.
- SupFig. S7 Correlation between maximum differences in metabolic gene expression and entropy.

#### Supporting tables

- SupTable S1 List of the metabolic genes used to estimate cell fluxomes.
- SupTable S2 List of the top19 variable metabolites.
- SupTable S3 Functional annotation clusters.
- SupTable S4 Lists of differentially expressed genes between clusters #0 and #1 of D4-expression data.
- SupTable S5 List of the metabolic modules selected for fluxome analysis.
- SupTable S6 Lists of genes within the clustered entropy profiles.
- SupTable S7 Gene set enrichment analysis: genes involved in biological processes or motif.

**SupFig. S1** Metabolic gene longitudinal expression. The mean expressions of the 530 metabolic genes were calculated at each collection day and are represented as a function of time. Although, we cannot assume monotonous variations between timepoints those are connected for visualization of dynamical patterns.

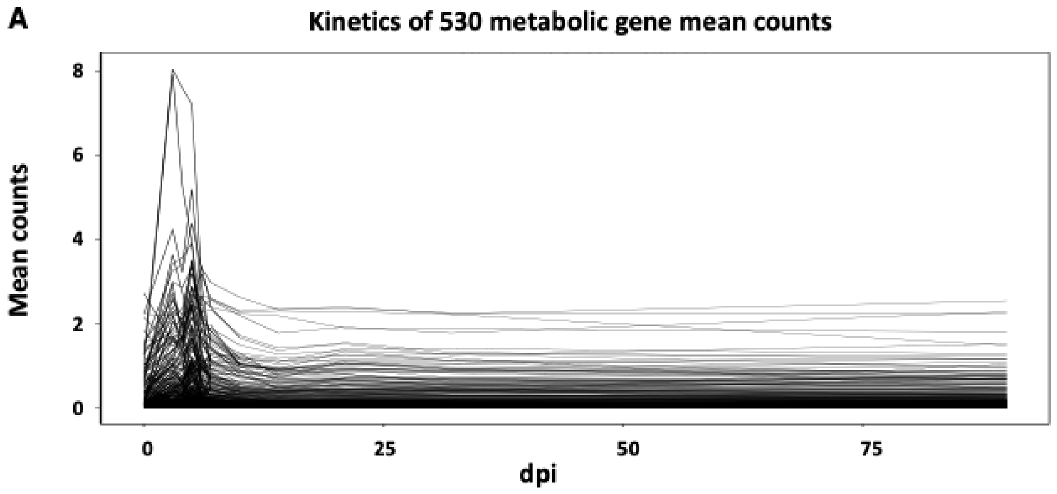

**SupFig. S2** Selection of flux modules with substantial variation. For each metabolic module, the distributions of flux values (x$10^3$ AU) at all dpi are shown left-hand and a box-plot of flux values for all cells is shown right-hand. The vertical bar on the left and the red dot on the right indicate the median and mean of cell flux values, respectively. The |CV| of flux values is indicated. (A) Examples of discarded modules with no variation (|CV| = 0). (B) Examples of discarded modules with little variation (0 < |CV| <$7.10^{-4}$). (C) Examples of retained modules with substantial flux variation (|CV| > $7.10^{-4}$). (D) Re-introduced M-114 module with substantial variation but a little |CV| ($2.10^{-4}$).

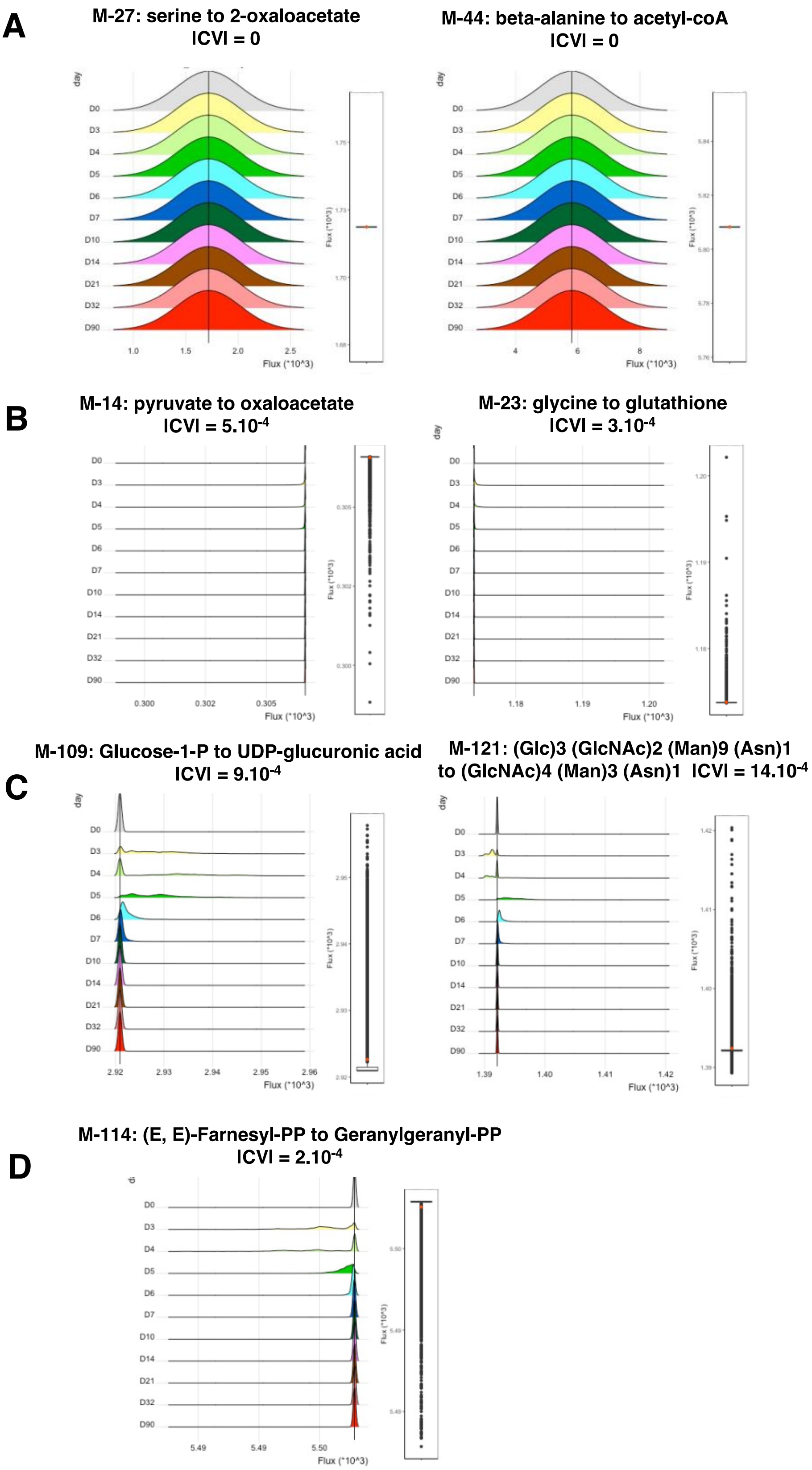

**SupFig. S3** Selection of modules with relevant flux values. (A) Heatmap of the flux values in all cells for the 47 modules selected for substantial flux variations. The heatmap is colored so that negative flux values appear in blue and positive flux values inferior to $10^{-3}$ AU in white. Cells in rows are colored according to the time of harvest with the same key as everywhere else in the manuscript. Modules in columns are colored according to their Super-Module family as in Alghamdi et al. [30]. (B, C) For each metabolic module, the distributions of flux values (x$10^{3}$ AU) at all dpi are shown left-hand and a box-plot of flux values for all cells is shown right-hand. The vertical bar on the left and the red dot on the right indicate the median and mean of flux values, respectively. The median of flux values is indicated. (B) Examples of discarded modules with very weak fluxes (median < $3.10^{-4}$ AU). (C) Examples of retained modules with weak fluxes but a significant fraction of cells that reach the $10^{-3}$ AU threshold (median > $3.10^{-4}$ AU).

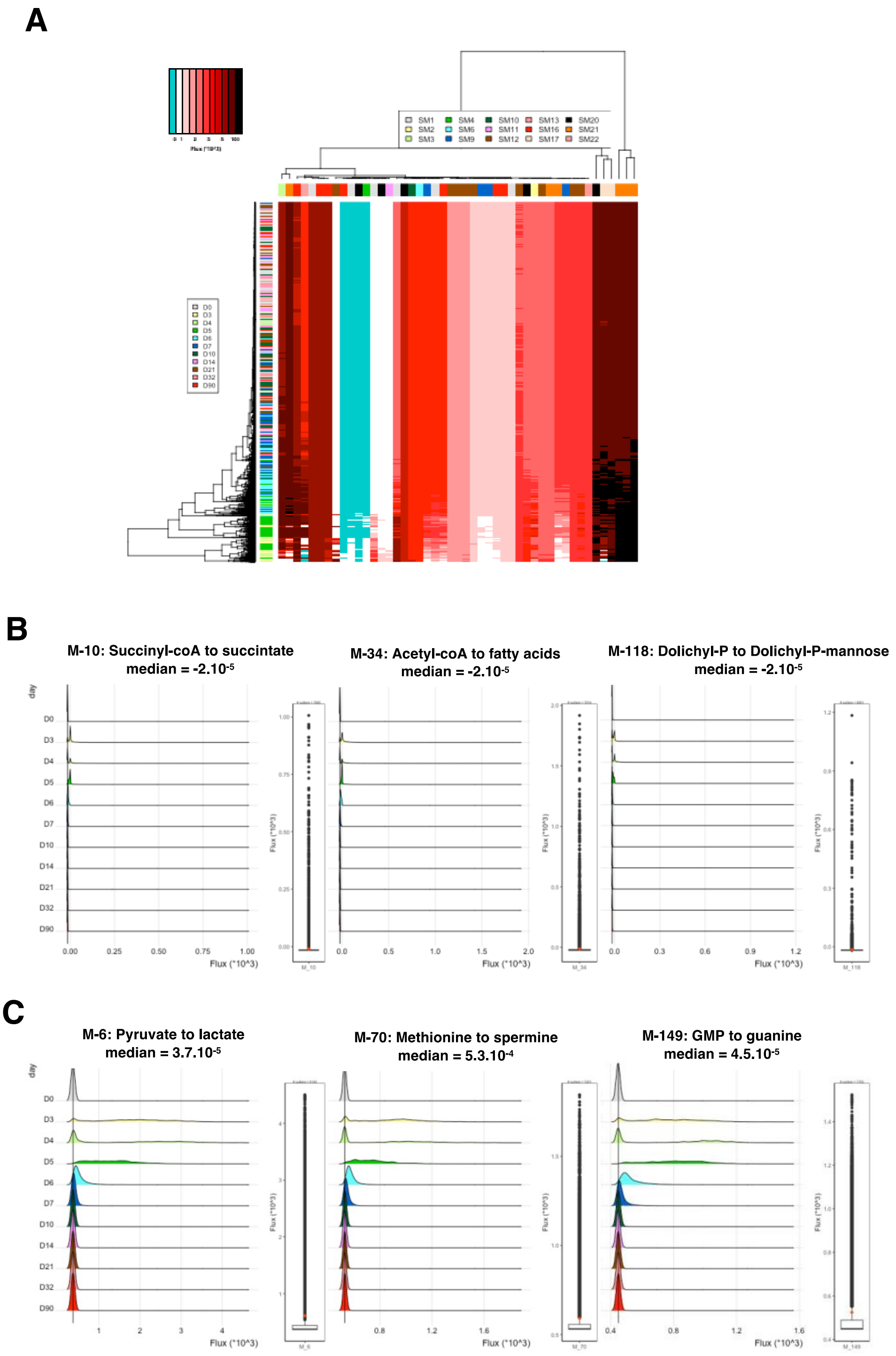

**SupFig. S4** Distribution of metabolite concentration_deltas. For each of the 70 metabolites for which scFEA returned concentration values, means were calculated at each dpi and the distribution of concentration_deltas, corresponding to the difference between the highest and lowest means, is depicted.

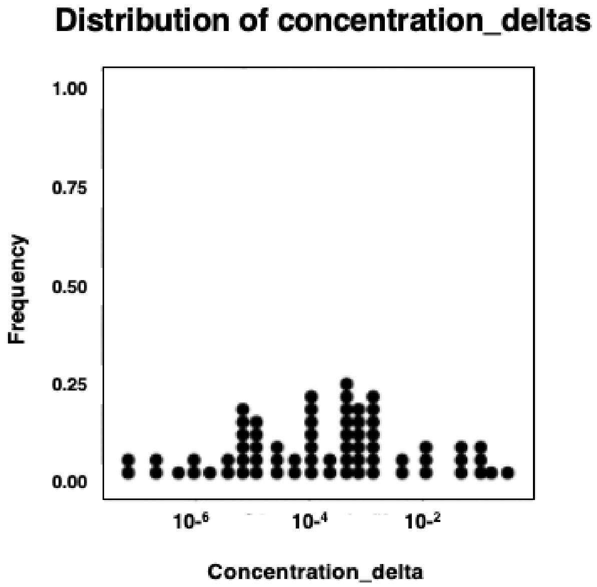

**SupFig. S5** Variations of the concentrations of the top19 variable metabolites. For each indicated metabolite, the deviations of minimal (left-hand) and maximal (right-hand) values of concentration from the median of all cell values are represented (AU). For metabolites showing a simple depletion or accumulation profile the time of the peak is indicated. For metabolites with biphasic or triphasic changes in concentrations, the times of all minima and maxima peaks are indicated.

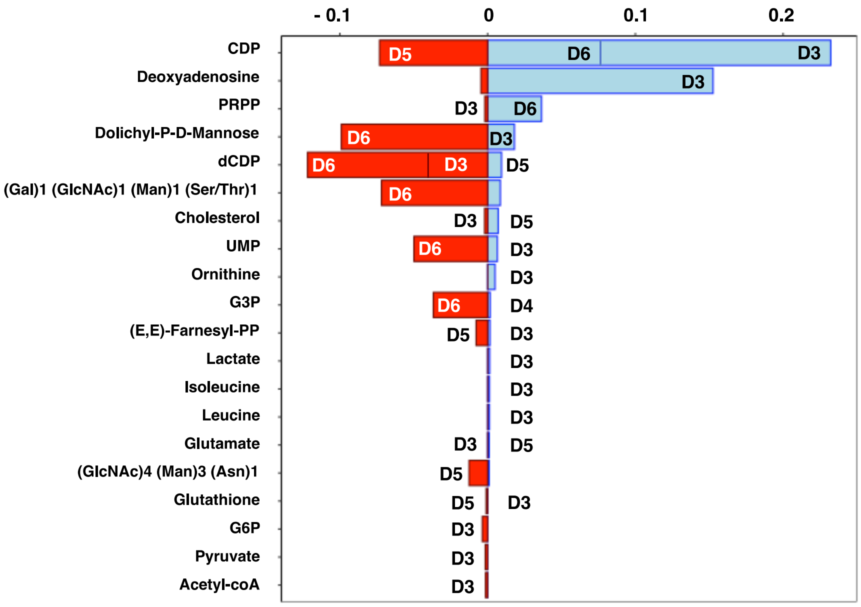

**SupFig. S6** Fluxome representation of all the 34 modules selected for analysis. For each metabolic module, the distributions of flux values (x$10^3$ AU) at all dpi are shown left-hand and a box-plot of flux values for all cells is shown right-hand. The vertical bar on the left and the red dot on the right indicate the median and mean of flux values, respectively. Names, as well as the initial and final compounds, of modules are indicated in the color of the corresponding Super-Module, as in Fig 3.

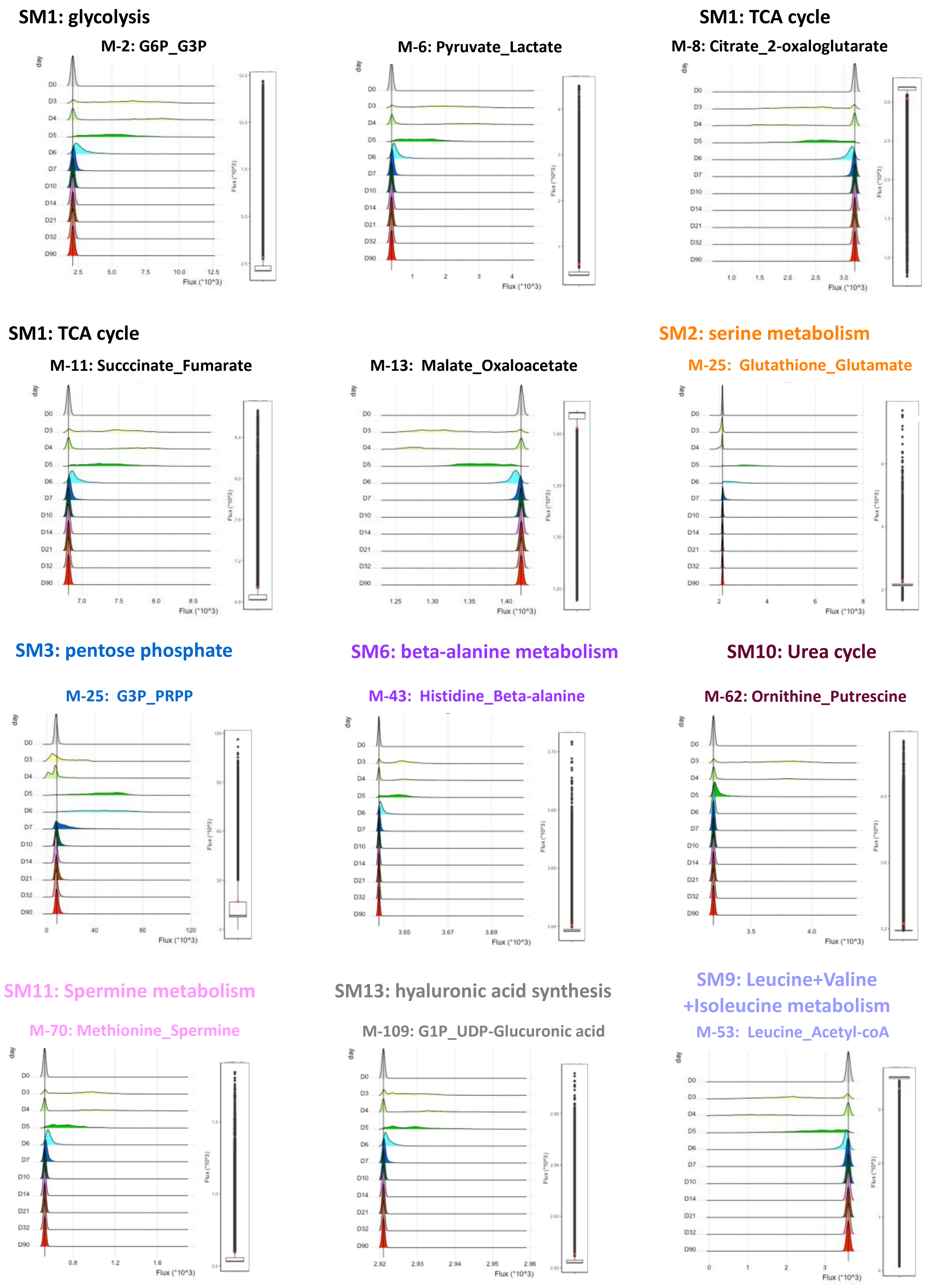

#### SM9: Leucine+Valine+Isoleucine metabolism

##### M-55: Isoleucine_Succinyl-coA

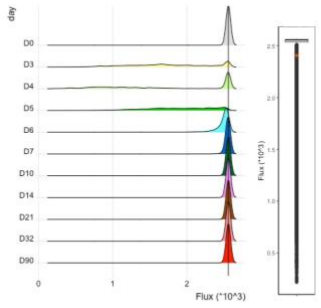


##### M-56: Isoleucine _Acetyl-coA

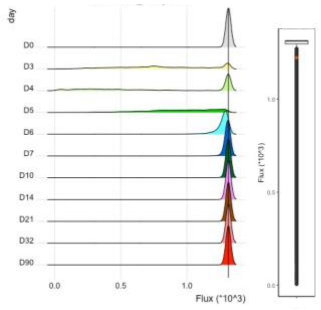


##### M-60: Lysine_Acetyl-coA

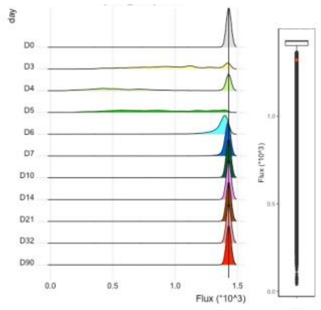


#### SM16: N-linked glycan synthesis

##### M-113: Acetyl-coA_E,E-Farnesyl-PP

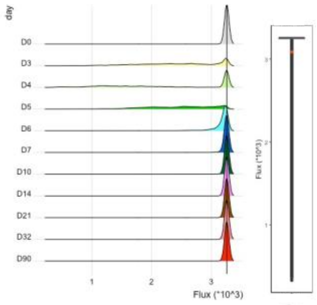


##### M-114: E,E-Farnesyl-PP _Geranylgeranyl-P

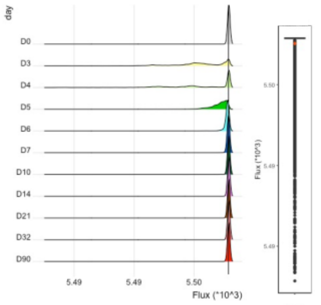


##### M-115: E,E-Farnesyl-PP _Farnesal

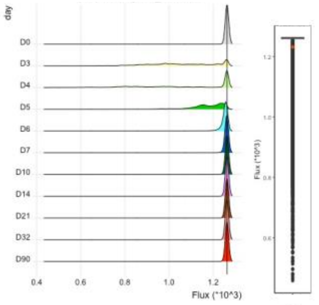


#### SM16: N-linked glycan synthesis

##### M-119: dolychil-P_ (GlcNAc)4 (Man)3 (Asn)1

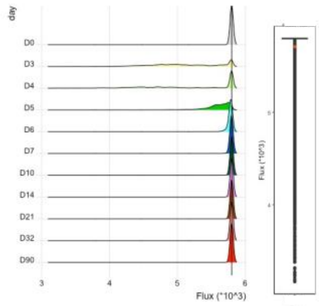


##### M-121: (Glc)3 (GlcNAc)2 (Man)9 (Asn)1 _(GlcNAc)4 (Man)3 (Asn)1

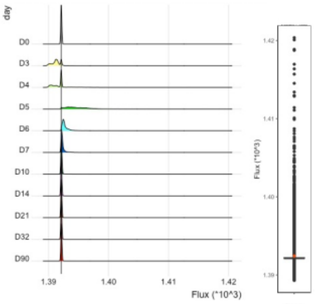


##### M-122: (GlcNAc)4 (Man)3 (Asn)1_(Gal)2 (GlcNAc)4 (LFuc)1 (Man)3 (Neu5Ac)2 (Asn)1

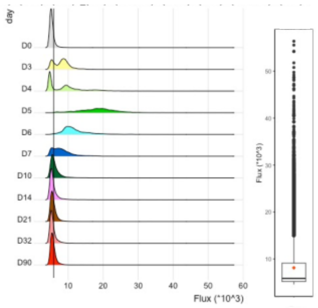


#### SM17: O-linked glycan synthesis

##### M-125: Dolichyl-P-Mannose + Prot-Serine_ _(Gal)1 (GlcNAc)1 (Man)1 (Ser/Thr)1

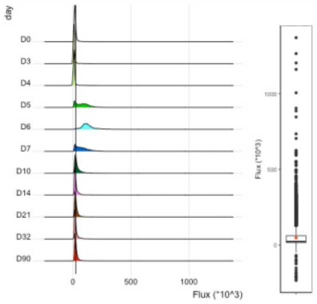


##### M-126: (Gal)1 (GlcNAc)1 (Man)1 (Ser/Thr)1_ (Gal)1 (GlcNAc)1 (Man)1 (Neu5Ac)1 (Ser/Thr)1

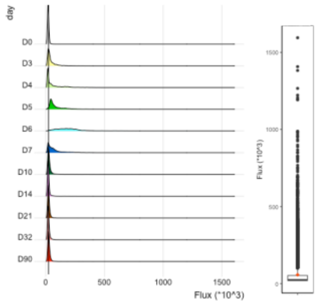


#### SM22: Cholesterol synthesis

##### M-167: E,E-Farnesyl-PP_cholesterol

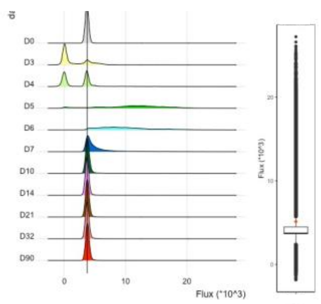

#### SM21: Pyrimidine synthesis

##### M-153: UMP_CDP

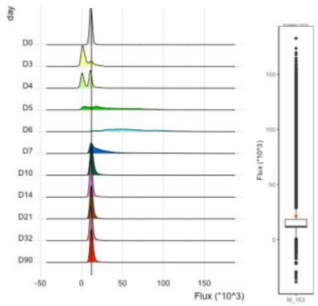


##### M-155: URP_CDP

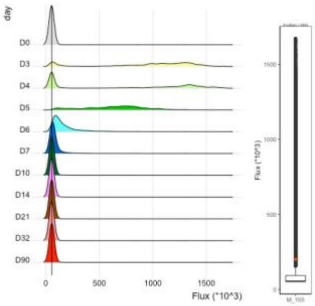


##### M-157: CDP_dCDP

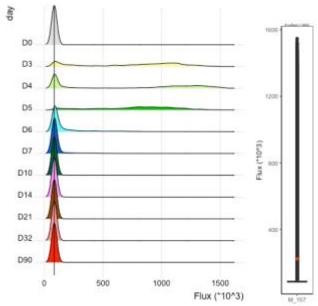


#### SM21: Pyrimidine synthesis

##### M-158: dCDP_Deoxycytidine

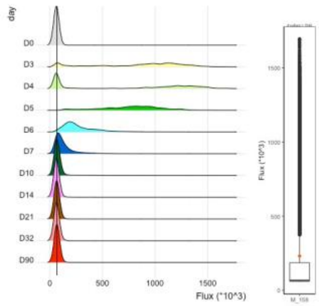


##### M-161: dCDP_dCTP

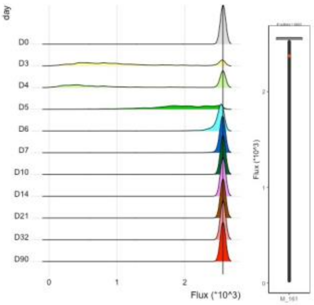


##### M-171: dCDP_dCMP

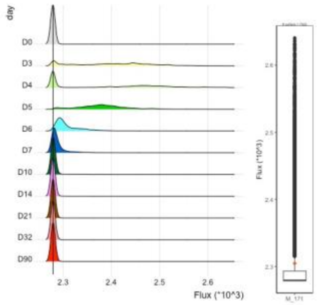


#### SM20: Purine synthesis

##### M-135: AICAR_IMP

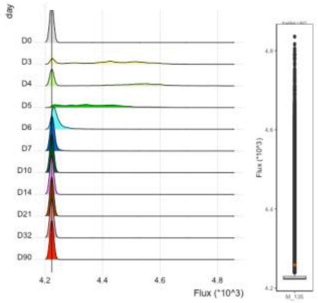


##### M-140: ADP_Deoxyadenosine

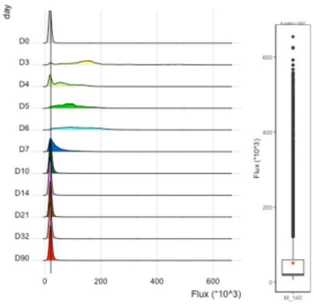


##### M-143: IMP_Hypoxanthine

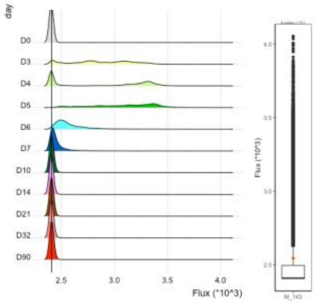


#### SM20: Purine synthesis

##### M-149: GMP_Guanine

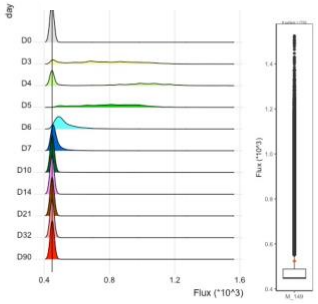

**SupFig. S7** Correlation between maximum differences in metabolic gene expression and entropy. For each of the 530 metabolic genes, the mean expression and entropy was calculated at each time point and the delta between maximal and minimal values for expression levels and entropies are plotted one against the other. The Pearson's correlation coefficient is indicated.

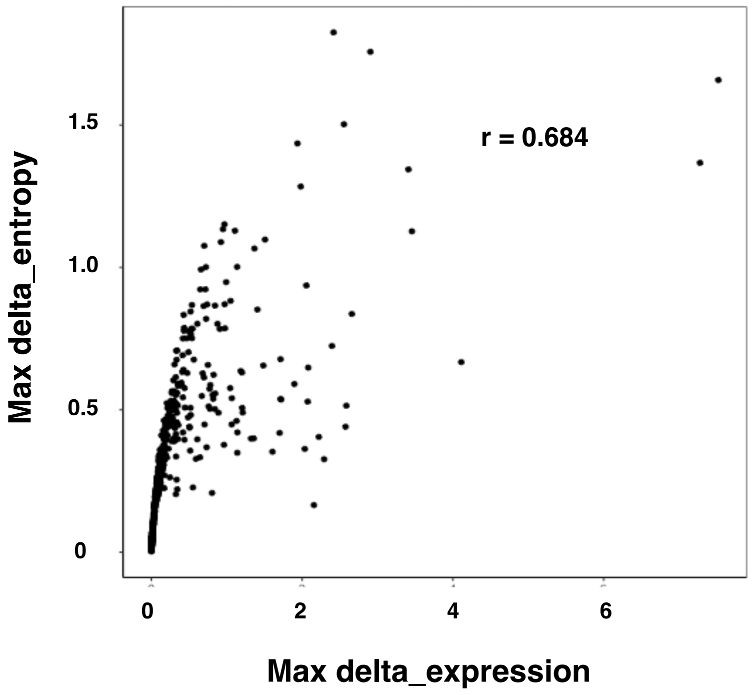

**SupTable S1.** List of the metabolic genes used to estimate cell fluxomes.
The origin, KEGG database release '101.4' or Alghamdi et al. [30], of genes is indicated.

#### Genes from N. Alghamdi et al. 2021

| | | | | | | |
|---|---|---|---|---|---|---|
| Aacs | Amdhd2 | Dhodh | Got1 | Mdh1 | Pgm2 | Slc6a6 |
| Acaa1a | Ampd1 | Dlat | Got2 | Mdh2 | Pgm3 | Slc7a1 |
| Acaa1b | Ampd2 | Dld | Gpi1 | Mecr | Phb | Slc7a5 |
| Acaa2 | Aoc2 | Dlst | Gss | Mgat1 | Phgdh | Slc7a6 |
| Acaca | Apip | Dnmt1 | Gsta4 | Mgat4a | Phykpl | Slc7a7 |
| Acacb | Aprt | Dnmt3a | Gstk1 | Mgat4b | Pipox | Smox |
| Acad8 | Asl | Dnmt3b | Gstm1 | Mgst2 | Pkm | Sms |
| Acadl | Asns | Dpm1 | Gstm4 | Mgst3 | Pmvk | Sqle |
| Acadm | Asrgl1 | Dse | Gstm5 | Mif | Pnp | Srd5a1 |
| Acads | Ass1 | Dtymk | Gsto1 | Minpp1 | Poglut1 | Srd5a3 |
| Acadsb | Atic | Ebp | Gstp1 | Mlycd | Pomgnt1 | Srm |
| Acadvl | Auh | Echdc1 | Gstp3 | Mogs | Pomt1 | St3gal2 |
| Acat1 | B3galt6 | Echs1 | Gstt2 | Mpst | Pomt2 | St3gal3 |
| Acat2 | B3gat3 | Ehhadh | Gstz1 | Mri1 | Ppat | St3gal6 |
| Acat3 | B3glct | Eno1 | Gys1 | Msmo1 | Ppt1 | St6gal1 |
| Acly | B4galt1 | Eno3 | H6pd | Mtap | Prps1 | Stt3a |
| Aco1 | B4galt3 | Enoph1 | Hadh | Mtr | Prps2 | Stt3b |
| Aco2 | B4galt7 | Entpd1 | Hadha | Mvd | Psat1 | Sucla2 |
| Acot1 | Bcat1 | Entpd5 | Hadhb | Mvk | Psph | Suclg1 |
| Acot2 | Bcat2 | Entpd6 | Hibadh | Naalad2 | Pycr1 | Suclg2 |
| Acot7 | Bckdha | Ext1 | Hibch | Nans | Pycr2 | Sult2b1 |
| Acot8 | Bckdhb | Ext2 | Hk1 | Nme1 | Pycrl | Taldo1 |
| Acox1 | Bpgm | Extl1 | Hk2 | Nme2 | Pygb | Tk1 |
| Acox3 | Cad | Extl2 | Hk3 | Nme3 | Rbks | Tk2 |
| Acsbg1 | Cant1 | Extl3 | Hmgcl | Nme4 | Rce1 | Tkt |
| Acsf3 | Carns1 | Fah | Hmgcr | Nme6 | Rdh10 | Tktl1 |
| Acsl1 | Cbs | Fasn | Hmgcs1 | Nme7 | Rimkla | Tm7sf2 |
| Acsl3 | Cdo1 | Fdft1 | Hprt | Nsdhl | Rnf2 | Tpi1 |
| Acsl4 | Chpf | Fdps | Hsd11b1 | Nt5c | Rpe | Tspan31 |
| Acsl5 | Chpf2 | Fh1 | Hsd17b10 | Nt5c2 | Rpia | Tst |
| Acsl6 | Chst10 | Fnta | Hsd17b12 | Nt5c3 | Rrm1 | Tyms |
| Ada | Chsy1 | Fntb | Hsd17b4 | Nt5c3b | Rrm2 | Uap1 |
| Adi1 | Clk1 | Fut4 | Hsd17b7 | Nt5e | Rrm2b | Uap1l1 |
| Adk | Cmas | Fut7 | Hykk | Nt5m | Sc5d | Uck1 |
| Adpgk | Cmpk1 | Fut8 | Icmt | Nudt16 | Scp2 | Uck2 |
| Adsl | Cmpk2 | G6pc3 | Idh1 | Nudt2 | Sdha | Uckl1 |
| Adss | Cndp2 | G6pdx | Idh2 | Nudt5 | Sdhb | Ugdh |
| Adssl1 | Comt | Galm | Idh3a | Oat | Shmt1 | Ugp2 |
| Ahcy | Coq2 | Gamt | Idh3b | Odc1 | Shmt2 | Umps |
| Ahcyl1 | Coq3 | Ganab | Idh3g | Ogdh | Slc15a4 | Upb1 |
| Ahcyl2 | Coq5 | Gapdh | Idi1 | Ogdhl | Slc16a1 | Upp1 |
| Ak2 | Coq6 | Gart | Impdh1 | Oplah | Slc16a3 | Uprt |
| Ak3 | Coq7 | Gatm | Impdh2 | Oxct1 | Slc16a4 | Xdh |
| Ak4 | Cpt1a | Gcat | Ivd | Oxsm | Slc17a7 | Xylt1 |
| Ak6 | Cpt2 | Gcdh | Kyat1 | Paics | Slc1a4 | Xylt2 |
| Ak7 | Cs | Gclc | Kyat3 | Pcca | Slc1a5 | Zmpste24 |
| Alas1 | Csgalnact2 | Gfpt1 | Lap3 | Pccb | Slc25a1 | |
| Aldh1b1 | Cth | Gfpt2 | Ldha | Pck2 | Slc25a10 | |
| Aldh2 | Ctps | Ggct | Ldhb | Pcx | Slc25a11 | |
| Aldh3a2 | Ctps2 | Ggps1 | Ldhd | Pcyox1 | Slc25a15 | |
| Aldh3b1 | Cyp11a1 | Ggt1 | Lss | Pdha1 | Slc25a29 | |
| Aldh5a1 | Cyp17a1 | Ggt5 | Man1a | Pdhb | Slc27a1 | |
| Aldh7a1 | Cyp2d22 | Ggt7 | Man1a2 | Pdss1 | Slc27a3 | |
| Aldh9a1 | Cyp2s1 | Gldc | Man1b1 | Pdss2 | Slc27a4 | |
| Aldoa | Cyp39a1 | Gls | Man1c1 | Pecr | Slc29a1 | |
| Aldoc | Cyp51 | Gls2 | Man2a1 | Pfas | Slc29a2 | |
| Alg10b | Dbt | Glud1 | Man2a2 | Pfkl | Slc2a1 | |
| Alg11 | Dck | Glul | Maoa | Pfkm | Slc2a3 | |
| Alg5 | Dctd | Gm5566 | Mat2a | Pfkp | Slc33a1 | |
| Alg6 | Dctpp1 | Gmps | Mat2b | Pgam1 | Slc36a1 | |
| Alg8 | Ddc | Gne | Mcat | Pgd | Slc38a1 | |
| Amacr | Dhcr24 | Gnpda1 | Mccc1 | Pgk1 | Slc38a2 | |
| Amd1 | Dhcr7 | Gnpda2 | Mccc2 | Pgls | Slc6a13 | |
| Amdhd1 | Dhdds | Gnpnat1 | Mcee | Pgm1 | Slc6a20b | |

#### Extra genes from KEGG database release '101.4', involved in glycolysis

Acss1
Acss2
Adh4
Akr1a1

#### Extra genes from KEGG database release '101.4', involved in oxydative phosphorylation

Atp6v1h
Atp5a1
Atp5b
Atp5c1
Atp5d
Atp5e
Atp5g1
Atp5g2
Atp5g3
Atp5h
Atp5j
Atp5j2
Atp5k
Atp5o
Atp5pb
Atp6v0a1
Atp6v0a2
Atp6v0b
Atp6v0c
Atp6v0d1
Atp6v0e
Atp6v1a
Atp6v1b2
Atp6v1c1
Atp6v1d
Atp6v1e1
Atp6v1f
Atp6v1g1
Atp6v1g2
Atp6v1g3
Cox10
Cox11
Cox15
Cox17
Cox4i1
Cox4i2
Cox5a
Cox5b
Cox6a1
Cox6a2
Cox6b1
Cox6b2
Cox6c
Cox7a1
Cox7a2
Cox7b
Cox7c
Cox8a
Ndufa1
Ndufa10
Ndufa11
Ndufa12
Ndufa13
Ndufa2
Ndufa3
Ndufa4
Ndufa5
Ndufa6
Ndufa7
Ndufa8
Ndufa9
Ndufab1
Ndufb10
Ndufb11
Ndufb2
Ndufb3
Ndufb4
Ndufb5
Ndufb6
Ndufb7
Ndufb8
Ndufb9
Ndufc1
Ndufc2
Ndufs1
Ndufs2
Ndufs3
Ndufs4
Ndufs5
Ndufs6
Ndufs7
Ndufs8
Ndufv1
Ndufv2
Ndufv3
Ppa1
Ppa2
Tcirg1
Uqcr10
Uqcr11
Uqcrb
Uqcrc1
Uqcrc2
Uqcrfs1
Uqcrh
Uqcrq

**SupTable S2.** List of the top19 variable metabolites.
The metabolites are grouped by kinetics of fluctuations and ordered in decreasing delta of concentrations. The corresponding Super-Modules are indicated, as well as the direction of fluctuations and the maximal delta of concentrations.

| Metabolite | Fluctuation | Super-Module (Jonction) | Concentration delta ($10^{-3}$ AU) |
|---|---|---|---|
| **Accumulated** | | | |
| Deoxyadenosine | +/+ | SM20 | 157.1 |
| Lactate | +/+ | SM1 | 1.425 |
| Isoleucine | +/+ | SM9 | 1.270 |
| Leucine | +/+ | SM9 | 1.235 |
| **Deprived** | | | |
| (Gal)1 (GlcNAc)1 (Man)1 (Ser/Thr)1 | -/- | SM17 | 80.46 |
| (GlcNAc)4 (Man)3 (Asn)1 | -/- | SM16 | 13.58 |
| G6P | -/- | SM1 | 3.653 |
| Pyruvate | -/- | SM1 | 1.615 |
| Acetyl-coA | -/- | SM1 (SM9) | 1.403 |
| **Deprived then accumulated** | | | |
| PRPP | -/+ | SM3 | 38.45 |
| Cholesterol | -/+ | SM22 | 9.204 |
| Glutamate | -/+ | SM2 | 1.082 |
| **Accumulated then deprived** | | | |
| Dolichyl-P-D-Mannose | +/- | SM17 | 117.1 |
| UMP | +/- | SM21 | 56.48 |
| G3P | +/- | SM1 (SM3) | 38.44 |
| 2-trans,6-trans-Farnesyl-PP | +/- | SM16 (SM22) | 9.408 |
| Glutathione | +/- | SM2 | 1.081 |
| **Fluctuating** | | | |
| dCDP | -/+/- | SM21 | 131.3 |
| CDP | +/-/+ | SM21 | 305.4 |

+/+ denotes metabolites accumulated in cells between D0 and D10
-/- denotes metabolites deprived from cells between D0 and D10
-/+ denotes metabolites deprived from cells between D0 and D4 and then transiently accumulated
+/- denotes metabolites accumulated in cells between D0 and D4 and then transiently deprived
-/+/- and +/-/+ denote fluctuating metabolites

**SupTable S3.** Functional annotation clusters. GO terms and enrichment scores (*ES) of the functional annotation clustering of the gene lists corresponding to clustered entropy profiles.

| 78/86 genes of immediate entropy cluster 1 | | |
|---|---|---|
| Cluster | ES* | GO terms |
| 1 | 14.86 | mitochondrion<br>mitochondrion inner membrane<br>membrane |
| 2 | 11.29 | mitochondrion inner membrane<br>mitochondrial ATP synthesis coupled proton transport<br>aerobic respiration<br>respiratory chain<br>mitochondrial respiratory chain complex I<br>mitochondrial respiratory chain complex I assembly<br>NADH dehydrogenase (ubiquinone) activity |
| 3 | 9.06 | 'de novo' XMP biosynthetic process<br>GMP biosynthetic process<br>purine nucleotide biosynthetic process<br>'de novo' IMP biosynthetic process<br>'de novo' AMP biosynthetic process |
| 4 | 6.06 | respiratory chain<br>mitochondrial respiratory chain complex III<br>mitochondrial electron transport, ubiquinol to cytochrome-c<br>cellular respiration<br>ubiquinol-cytochrome-c reductase activity |
| 5 | 4.91 | transferase activity<br>nucleoside diphosphate kinase activity<br>ATP binding<br>nucleotide binding<br>UTP biosynthetic process<br>CTP biosynthetic process<br>GTP biosynthetic process<br>phosphorylation<br>kinase activity<br>nucleoside diphosphate phosphorylation<br>nucleoside metabolic process |
| 6 | 4.57 | glycolytic process<br>carbohydrate metabolic process<br>glucogenesis |
| 7 | 3.72 | ATP synthesis coupled proton transport<br>ATP biosynthetic process<br>mitochondrial proton-transporting ATP synthase complex<br>hydrogen ion transmembrane transport<br>ATP biometabolic process<br>proton-transporting ATP synthase activity, rotational mechanism<br>ion transport |
| 8 | 2.96 | tetrahydrofolate metabolic process<br>translation repressor activity, nucleic acid binding<br>tetrahydrofolate interconversion |
| 9 | 2.95 | ATP synthesis coupled proton transport<br>mit. proton-transporting ATP synthase complex, coupling factor F(o)<br>proton-transporting ATP synthase complex, coupling factor F(o)<br>hydrogen ion transmembrane transporter activity<br>ion transport |
| 10 | 1.58 | response to drugs<br>methylation<br>methyl transferase activity |

| 107/444 genes of later entropy cluster 2 | | |
|---|---|---|
| Cluster | ES* | GO terms |
| 1 | 31.23 | respiratory chain<br>aerobic respiratin<br>mitochondrial respiratory chain complex I<br>mitochondrial respiratory chain complex I assembly |
| 2 | 7.51 | long-chain fatty acid-coA ligase activity<br>arachidonate-coA igase activity<br>fatty acid transport |
| 3 | 3.51 | acetyl-coA biosynthetic process from pyruvate<br>pyruvate dehydrogenase (NAD+) activity<br>mitochondrial pyruvate dehydrogenase complex<br>pyruvate dehydrogenase complex<br>mitochondrial acetyl-coA biosynthetic process from pyruvate |
| 4 | 3.48 | dUMP catabolic process<br>dTMP catabolic process<br>UMP catabolic process<br>dCMP catabolic process<br>CMP catabolic process |
| 5 | 3.42 | allantoin metabolic process<br>IMP catabolic process<br>dGMP catabolic process |
| 6 | 2.76 | nucleoside-diphosphate activity<br>guanosine-diphosphate activity<br>uridine-diphosphate activity<br>adenosine-diphosphate activity<br>nucleoside-diphosphate catabolic process |
| 7 | 2.69 | intramolecular transferase activity, phosphotransferases<br>phosphoglucomutase activity<br>organic substance metabolic process |
| 8 | 2.64 | 6-phosphofructokinase complex<br>fructose 1,6-biphosphate metabolic process<br>6-phosphofructokinase activity<br>glycolytic process through fructose-6-phosphate<br>fructose-6-phosphate binding |
| 9 | 2.63 | glucoronosyl-N-acetylgalactosaminyl-proteoglycan 4-beta-N-acetylgalactosaminyltransferase activity<br>N-acetylgalactosaminyl-proteoglycan 3-glucoronosyltransferase activity<br>acetylgalactosaminyltransferase activity |
| 10 | 2.4 | palmitoyl-coA hydrolase activity<br>myristoyl-coA hydrolase activity<br>acetyl-coA hydrolase activity |
| 11 | 2.3 | AMP catabolic process<br>deoxyadenosine catabolic process<br>adenosine catabolic process<br>dAMP catabolic process |
| 12 | 2.15 | S-adenosylmethionine cycle<br>adenosylhomocysteinase activity<br>one-carbon metabolic process |
| 13 | 2.04 | beta-N-acetylglucosaminylglycopeptide beta-1,4-galactosyltransferase activity<br>glycosylation<br>galactosyltransferase activity |
| 14 | 1.82 | peptidyltransferase activity<br>glutathione hydrolase activity<br>glutathione catabolic process<br>response to tumor necrosis factor<br>response to lipopolysaccharide |
| 15 | 0.14 | heterochromatin<br>chromatin binding<br>negative regulation of transcription from RNA polymerase II promoter |

**SupTable S4.** Lists of differentially expressed genes between clusters #0 and #1 of D4-expression data.
Genes with an average log2(FC) > 0.13 were selected and ranked by decreasing FC.

| Up-regulated in Cluster #0 | | | | | | Up-regulated in Cluster #1 | | |
|---|---|---|---|---|---|---|---|---|
| Gene | Average log2(FC) | Adj. p_value<br>* 0 for < e-300 | Gene | Average log2(FC) | Adj. p_value | Gene | Average log2(FC) | Adj. p_value |
| Gzma | 1.48 | 2.54 e-260 | Vim | 0.25 | 0 | Actb | 1.18 | 0 |
| Ccl4 | 0.97 | 4.37 e-20 | S100a11 | 0.24 | 0 | Pim1 | 1.01 | 5.19 e-206 |
| S100a6 | 0.91 | 0 | Gltp | 0.24 | 0 | Slc39a1 | 0.97 | 0.01 |
| Mt1 | 0.82 | 0 | Aurka | 0.24 | 0 | Abhd2 | 0.87 | 3.70 e-293 |
| Ccl3 | 0.78 | 5.66 e-23 | Tnfrsf4 | 0.24 | 3.00 e-134 | Alkbh5 | 0.84 | 6.44 e-220 |
| Ifng | 0.76 | 3.61 e-162 | Gmfg | 0.23 | 0 | Wtap | 0.83 | 1.26 e-90 |
| X1500009L16Rik | 0.71 | 4.70 e-216 | S1pr4 | 0.23 | 0 | Tubb5 | 0.79 | 0 |
| Mt2 | 0.65 | 1.77 e-291 | Ifi27 | 0.23 | 0 | Kdelr2 | 0.77 | 0 |
| Crip1 | 0.64 | 0 | Chchd10 | 0.23 | 0 | Pfas | 0.64 | 1.15 e-104 |
| Malat1 | 0.63 | 0 | Phf11b | 0.23 | 0 | Sephs2 | 0.62 | 0 |
| Gzmb | 0.61 | 0 | Zbp1 | 0.22 | 0 | Gnb1 | 0.62 | 0 |
| Nkg7 | 0.58 | 0 | Sh3bgrl3 | 0.22 | 0 | Birc6 | 0.61 | 2.36 e-120 |
| Ifi27l2a | 0.58 | 0 | H1f5 | 0.22 | 0 | Timmdc1 | 0.56 | 1.33 e-84 |
| Mt3 | 0.57 | 0 | Reep5 | 0.21 | 0 | Nup98 | 0.51 | 1.29 e-95 |
| Hist1h2ap | 0.54 | 0 | Gm19585 | 0.21 | 0 | Kmt2d | 0.51 | 0 |
| Lgals3 | 0.53 | 3.29 e-132 | Ccr7 | 0.21 | 3.00 e-262 | Zc3hav1 | 0.49 | 1.22 8e-109 |
| Trbc1 | 0.53 | 0 | Marcksl1 | 0.20 | 0 | Itpripl1 | 0.48 | 6.71 e-174 |
| Batf3 | 0.52 | 0 | Id2 | 0.20 | 0 | Actr2 | 0.48 | 0 |
| S100a4 | 0.52 | 5.82 e-149 | Timm10 | 0.20 | 0 | Polh | 0.45 | 9.09 e-109 |
| Tomm5 | 0.51 | 0 | Ddx21 | 0.20 | 0 | Rrn3 | 0.44 | 1.43 e-130 |
| Ccl5 | 0.49 | 5.15 e-47 | Chchd4 | 0.20 | 0 | Dpysl2 | 0.43 | 7.75 e-134 |
| AA467197 | 0.48 | 8.64 e-73 | Bcl2a1b | 0.20 | 0 | Cdk6 | 0.43 | 0 |
| Lgals1 | 0.46 | 0 | Rsrp1 | 0.20 | 0 | Pan3 | 0.42 | 1.04 e-189 |
| AW112010 | 0.46 | 0 | Zfas1 | 0.19 | 0 | Ythdf2 | 0.40 | 0 |
| S100a10 | 0.46 | 0 | Smap2 | 0.19 | 0 | St13 | 0.39 | 0 |
| Anxa2 | 0.45 | 0 | Top2a | 0.19 | 0 | Yme1l1 | 0.38 | 1.76 e-230 |
| Itgb7 | 0.43 | 2.87 e-252 | Jund | 0.19 | 0 | Arglu1 | 0.36 | 0 |
| Ms4a4b | 0.40 | 0 | Npc2 | 0.19 | 0 | Mcmbp | 0.36 | 4.81 e-274 |
| Klrd1 | 0.40 | 5.59 e-71 | Ifitm3 | 0.19 | 1.12 e-179 | Gm2a | 0.33 | 2.06 e-177 |
| Trgc2 | 0.38 | 1.90 e-23 | Il2ra | 0.17 | 0 | Otub1 | 0.33 | 0 |
| Isg15 | 0.38 | 0 | Batf | 0.17 | 0 | Gsk3b | 0.32 | 6.18 e-171 |
| Ly6c2 | 0.37 | 0 | Glipr2 | 0.17 | 0 | Kdm6b | 0.31 | 5.71 e-40 |
| Klf2 | 0.35 | 0 | Glrx | 0.16 | 1.20 e-294 | Golt1b | 0.31 | 1.30 e-199 |
| Prf1 | 0.35 | 0 | Cdk2ap2 | 0.16 | 0 | Gtf2a1 | 0.28 | 2.05 e-215 |
| Serpinb6b | 0.34 | 0 | Fdps | 0.16 | 0 | Khsrp | 0.25 | 2.84 e-224 |
| H1f4 | 0.33 | 0 | Tspo | 0.16 | 0 | Srrm2 | 0.22 | 0 |
| Odc1 | 0.33 | 0 | Mki67 | 0.16 | 0 | Tmf1 | 0.21 | 0 |
| Btg1 | 0.33 | 1.04 e-103 | Srgn | 0.16 | 0 | Picalm | 0.20 | 0 |
| Gnai2 | 0.32 | 0 | Nefh | 0.16 | 1.26 e-243 | Klf13 | 0.18 | 0 |
| Cenpf | 0.32 | 0 | Nip7 | 0.16 | 0 | Furin | 0.18 | 5.36 e-140 |
| Ddt | 0.30 | 0 | Ech1 | 0.15 | 0 | B4galt1 | 0.18 | 0 |
| Bcl2a1d | 0.29 | 0 | Pten | 0.15 | 0 | Tacc1 | 0.17 | 5.88 e-283 |
| Emp3 | 0.29 | 0 | Junb | 0.15 | 0 | Hnrnpul1 | 0.16 | 0 |
| Nop58 | 0.29 | 0 | Cd160 | 0.15 | 0 | Hsf2 | 0.16 | 4.32 e-75 |
| Nfkbia | 0.29 | 0 | Fbxo5 | 0.15 | 0 | Lyrm4 | 0.15 | 4.73 e-213 |
| Cotl1 | 0.29 | 0 | Fyb | 0.15 | 0 | Rsl24d1 | 0.15 | 0 |
| Acadl | 0.28 | 0 | Cd8b1 | 0.15 | 0 | Map3k1 | 0.15 | 0 |
| Nolc1 | 0.27 | 0 | Timm8a1 | 0.14 | 0 | | | |
| Pycard | 0.27 | 0 | Incenp | 0.14 | 0 | | | |
| Desi2 | 0.27 | 0 | Saraf | 0.14 | 0 | | | |
| Rasgrp2 | 0.27 | 0 | Tuba1c | 0.13 | 0 | | | |
| Cmtm7 | 0.26 | 0 | H2ax | 0.13 | 0 | | | |
| Epsti1 | 0.26 | 0 | Hadh | 0.13 | 0 | | | |
| Flna | 0.26 | 0 | Septin1 | 0.13 | 0 | | | |
| Ctla2a | 0.26 | 0 | Rras2 | 0.13 | 0 | | | |
| Cenpe | 0.25 | 0 | | | | | | |

**SupTable S5.** List of the metabolic modules selected for fluxome analysis.
The modules are classified by Super-Modules as defined by Alghamdi et al. [30]. Initial and final compounds, as well as mean flux values and the direction of flux variation, are indicated.

| Module | From | To | Supermodule | Direction | Mean flux value ($10^{-3}$ AU) |
|---|---|---|---|---|---|
| M-2 | Glucose-6P (G6P) | Glyceraldehyde-3P (G3P) | SM1: Glycolysis | Up | 2.8 |
| M-6 | Pyruvate | Lactate | | Up | 0.6 |
| M-8 | Citrate | 2-Oxoglutarate | SM1: TCA cycle | Down | 3.0 |
| M-11 | Succinate | Fumarate | | Up | 6.9 |
| M-13 | Malate | Oxaloacetate | | Down | 1.4 |
| M-25 | Glutathione | Glutamate | SM2: Serine metabolism | Down, then up | 2.3 |
| M-33 | Glyceraldehyde-3P (G3P) | 5-Phosphoribose-2P (PRPP) | SM3: Pentose Phosphate | Down, then up | 16.9 |
| M-43 | Histidine | Beta-alanine | SM6: Beta-alanine metabolism | Up | 3.6 |
| M-53 | Leucine | Acetyl-coA | SM9: Leucine+Valine+Isoleucine | Down | 3.4 |
| M-55 | Isoleucine | Succinyl-coA | | Down | 2.4 |
| M-56 | Isoleucine | Acetyl-coA | | Down | 1.2 |
| M-60 | Lysine | Acetyl-coA | | Down | 1.3 |
| M-62 | Ornithine | Putrescine | SM10: Urea cycle | Up | 3.2 |
| M-70 | Methionine | Spermine | SM11: Spermine metabolism | Up | 0.6 |
| M-109 | Glucose-1P (G1P) | UDP-glucuronic acid | SM13: Hyaluronic acid synthesis | Up | 2.9 |
| M-113 | Acetyl-coA | 2-trans,6-trans-Farnesyl-PP | SM16: N-linked glycan synthesis | Down | 3.1 |
| M-114 | 2-trans,6-trans-Farnesyl-PP | Geranylgeranyl-PP | | Down | 5.5 |
| M-115 | 2-trans,6-trans-Farnesyl-PP | Farnesal | | Down | 1.2 |
| M-119 | Dolichyl-Phosphate | (GlcNAc)4 (Man)3 (Asn)1 | | Down | 5.7 |
| M-121 | (Glc)3 (GlcNAc)2 (Man)9 (Asn)1 | (GlcNAc)4 (Man)3 (Asn)1 | | Down, then up | 1.4 |
| M-122 | (GlcNAc)4 (Man)3 (Asn)1 | (Gal)2 (GlcNAc)4 (LFuc)1 (Man)3 (Neu5Ac)2 (Asn)1 | | Up | 8.1 |
| M-125 | Dolichyl-Phosphate-D-mannose + Protein serine | (Gal)1 (GlcNAc)1 (Man)1 (Ser/Thr)1 | SM17: O-linked glycan synthesis | Down, then up | 47.1 |
| M-126 | (Gal)1 (GlcNAc)1 (Man)1 (Ser/Thr)1 | (Gal)1 (GlcNAc)1 (Man)1 (Neu5Ac)1 (Ser/Thr)1 | | Up | 58.2 |
| M-135 | 5'-Phospho-ribosyl-5-amino-4-imidazole carboxamide (AICAR) | IMP | SM20: Purine synthesis | Up | 4.3 |
| M-140 | ADP | Deoxyadenosine | | Up | 51.0 |
| M-143 | IMP | Hypoxanthine | | Up | 2.5 |
| M-149 | GMP | Guanine | | Up | 0.5 |
| M-153 | UMP | CDP | SM21: Pyrimidne synthesis | Down, then up | 21.4 |
| M-155 | UTP | CDP | | Up | 201.0 |
| M-157 | CDP | dCDP | | Up | 223.0 |
| M-158 | dCDP | Deoxycytidine | | Up | 232.0 |
| M-161 | dCDP | dCTP | | Down | 2.4 |
| M-171 | dCDP | dCMP | | Up | 2.3 |
| M-167 | 2-trans,6-trans-Farnesyl-PP | Cholesterol | SM22: Steroid hormone synthesis | Down, then up | 5.1 |

**SupTable S6.** Lists of genes within the clustered entropy profiles.

| Cluster 1 | | Cluster 2 | | | | | | |
|---|---|---|---|---|---|---|---|---|
| Acadl | Psat1 | Aacs | Asl | Cox6b1 | Gfpt2 | Mat2b | Ogdhl | Slc38a2 |
| Acot7 | Psph | Acaa1a | Asrgl1 | Cox6b2 | Ggct | Mcat | Oplah | Slc6a13 |
| Adk | Pycrl | Acaa1b | Ass1 | Cox6c | Ggps1 | Mccc1 | Oxct1 | Slc6a20b |
| Adsl | Rrm1 | Acaa2 | Atp5c1 | Cox7a1 | Ggt1 | Mccc2 | Oxsm | Slc6a6 |
| Ak2 | Rrm2 | Acaca | Atp5d | Cox7a2 | Ggt5 | Mcee | Pcca | Slc7a1 |
| Ak6 | Sdhb | Acacb | Atp5e | Cox7b | Ggt7 | Mdh1 | Pccb | Slc7a5 |
| Aldoa | Shmt1 | Acad8 | Atp5g2 | Cox7c | Gldc | Mecr | Pck2 | Slc7a6 |
| Alg8 | Shmt2 | Acadm | Atp5h | Cox8a | Gls | Mgat1 | Pcx | Slc7a7 |
| Asns | Slc16a1 | Acads | Atp5j | Cpt1a | Gls2 | Mgat4a | Pcyox1 | Smox |
| Atic | Slc29a1 | Acadsb | Atp5j2 | Cpt2 | Glud1 | Mgat4b | Pdha1 | Sms |
| Atp5a1 | Srm | Acadvl | Atp5k | Csgalnact2 | Glul | Mgst2 | Pdhb | Sqle |
| Atp5b | Tkt | Acat1 | Atp5o | Cth | Gm5566 | Mgst3 | Pdss1 | Srd5a1 |
| Atp5g1 | Tpi1 | Acat2 | Atp6v0a1 | Ctps2 | Gmps | Minpp1 | Pdss2 | Srd5a3 |
| Atp5g3 | Tyms | Acat3 | Atp6v0a2 | Cyp11a1 | Gne | Mlycd | Pecr | St3gal2 |
| Atp5pb | Uck2 | Acly | Atp6v0b | Cyp17a1 | Gnpda1 | Mpst | Pfkl | St3gal3 |
| Bcat1 | Umps | Aco1 | Atp6v0c | Cyp2d22 | Gnpda2 | Mri1 | Pfkm | St3gal6 |
| Cad | Uqcr10 | Aco2 | Atp6v0d1 | Cyp2s1 | Gnpnat1 | Msmo1 | Pfkp | St6gal1 |
| Comt | Uqcr11 | Acot1 | Atp6v0e | Cyp39a1 | Gpi1 | Mtr | Pgls | Stt3a |
| Coq7 | Uqcrfs1 | Acot2 | Atp6v1a | Dbt | Gss | Mvd | Pgm1 | Stt3b |
| Cs | Uqcrq | Acot8 | Atp6v1b2 | Dck | Gsta4 | Mvk | Pgm2 | Sucla2 |
| Ctps | | Acox1 | Atp6v1c1 | Dctd | Gstk1 | Naalad2 | Pgm3 | Suclg1 |
| Cyp51 | | Acox3 | Atp6v1d | Ddc | Gstm1 | Nans | Phykpl | Suclg2 |
| Dctpp1 | | Acsbg1 | Atp6v1e1 | Dhcr24 | Gstm4 | Ndufa1 | Pipox | Sult2b1 |
| Dnmt1 | | Acsf3 | Atp6v1f | Dhcr7 | Gstm5 | Ndufa10 | Pmvk | Taldo1 |
| Dtymk | | Acsl1 | Atp6v1g1 | Dhdds | Gsto1 | Ndufa11 | Pnp | Tcirg1 |
| Enoph1 | | Acsl3 | Atp6v1g2 | Dhodh | Gstp1 | Ndufa13 | Poglut1 | Tk1 |
| Fasn | | Acsl4 | Atp6v1g3 | Dlat | Gstp3 | Ndufa2 | Pomgnt1 | Tk2 |
| Fdps | | Acsl5 | Atp6v1h | Dld | Gstz1 | Ndufa3 | Pomt1 | Tktl1 |
| Fh1 | | Acsl6 | Auh | Dlst | Gys1 | Ndufa4 | Pomt2 | Tm7sf2 |
| Got1 | | Acss1 | B3galt6 | Dnmt3a | H6pd | Ndufa6 | Ppa2 | Tspan31 |
| Got2 | | Acss2 | B3gat3 | Dnmt3b | Hadh | Ndufa7 | Ppt1 | Tst |
| Gstt2 | | Ada | B3glct | Dpm1 | Hadha | Ndufa8 | Prps2 | Uap1 |
| Hk2 | | Adh4 | B4galt1 | Dse | Hadhb | Ndufa9 | Pycr1 | Uap1l1 |
| Idh3a | | Adi1 | B4galt3 | Ebp | Hibadh | Ndufb10 | Pycr2 | Uck1 |
| Idi1 | | Adpgk | B4galt7 | Echdc1 | Hibch | Ndufb11 | Pygb | Uckl1 |
| Impdh2 | | Adss | Bcat2 | Echs1 | Hk1 | Ndufb3 | Rbks | Ugdh |
| Lap3 | | Adssl1 | Bckdha | Ehhadh | Hk3 | Ndufb5 | Rce1 | Ugp2 |
| Ldha | | Ahcy | Bckdhb | Eno1 | Hmgcl | Ndufb6 | Rdh10 | Upb1 |
| Mat2a | | Ahcyl1 | Bpgm | Eno3 | Hmgcr | Ndufb7 | Rimkla | **Upp1** |
| Mdh2 | | Ahcyl2 | Cant1 | Entpd1 | Hmgcs1 | Ndufb9 | Rnf2 | Uprt |
| Mif | | Ak3 | Carns1 | Entpd5 | Hprt | Ndufs1 | Rpe | Uqcrb |
| Mogs | | Ak4 | Cbs | Entpd6 | Hsd11b1 | Ndufs2 | Rpia | Uqcrc1 |
| Mtap | | Ak7 | Cdo1 | Ext1 | Hsd17b10 | Ndufs3 | Rrm2b | Uqcrc2 |
| Ndufa12 | | Akr1a1 | Chpf | Ext2 | Hsd17b12 | Ndufs4 | Sc5d | Uqcrh |
| Ndufa5 | | Alas1 | Chpf2 | Extl1 | Hsd17b4 | Ndufs5 | Scp2 | Xdh |
| Ndufab1 | | Aldh1b1 | Chst10 | Extl2 | Hsd17b7 | Ndufs6 | Sdha | Xylt1 |
| Ndufb2 | | Aldh2 | Chsy1 | Extl3 | Hykk | Ndufs7 | Slc15a4 | Xylt2 |
| Ndufb4 | | Aldh3a2 | Clk1 | Fah | Icmt | Ndufs8 | Slc16a3 | Zmpste24 |
| Ndufb8 | | Aldh3b1 | Cmas | Fdft1 | Idh1 | Ndufv1 | Slc16a4 | |
| Ndufc1 | | Aldh5a1 | Cmpk1 | Fnta | Idh2 | Ndufv2 | Slc17a7 | |
| Ndufc2 | | Aldh7a1 | Cmpk2 | Fntb | Idh3b | Ndufv3 | Slc1a4 | |
| Nme1 | | Aldh9a1 | Cndp2 | Fut4 | Idh3g | Nme3 | Slc1a5 | |
| Nme2 | | Aldoc | Coq2 | Fut7 | Impdh1 | Nme4 | Slc25a1 | |
| Nme6 | | Alg10b | Coq3 | Fut8 | Ivd | Nme7 | Slc25a10 | |
| Odc1 | | Alg11 | Coq5 | G6pc3 | Kyat1 | Nsdhl | Slc25a11 | |
| Paics | | Alg5 | Coq6 | G6pdx | Kyat3 | Nt5c | Slc25a15 | |
| Pfas | | Alg6 | Cox10 | Galm | Ldhb | Nt5c2 | Slc25a29 | |
| Pgam1 | | Amacr | Cox11 | Gamt | Ldhd | Nt5c3 | Slc27a1 | |
| Pgd | | Amd1 | Cox15 | Ganab | Lss | Nt5c3b | Slc27a3 | |
| Pgk1 | | Amdhd1 | Cox17 | Gapdh | Man1a | Nt5e | Slc27a4 | |
| Phb | | Amdhd2 | Cox4i1 | Gart | Man1a2 | Nt5m | Slc29a2 | |
| Phgdh | | Ampd1 | Cox4i2 | Gatm | Man1b1 | Nudt16 | Slc2a1 | |
| Pkm | | Ampd2 | Cox5a | Gcat | Man1c1 | Nudt2 | Slc2a3 | |
| Ppa1 | | Aoc2 | Cox5b | Gcdh | Man2a1 | Nudt5 | Slc33a1 | |
| Ppat | | Apip | Cox6a1 | Gclc | Man2a2 | Oat | Slc36a1 | |
| Prps1 | | Aprt | Cox6a2 | Gfpt1 | Maoa | Ogdh | Slc38a1 | |

**SupTable S7.** Gene set enrichment analysis: genes involved in biological processes or motif. The genes are ordered in decreasing fold-change of expression between both clusters.

| Cluster # 0 enriched biological processes and associated genes | | | |
|---|---|---|---|
| Response to stimulus | Response to stress | Response to external biotic stimulus | Response to other organisms |
| GzmA | | GzmA | GzmA |
| Ccl4 | Ccl4 | Ccl4 | Ccl4 |
| Mt1 | Mt1 | | |
| Ccl3 | Ccl3 | Ccl3 | Ccl3 |
| Ifng | Ifng | Ifng | Ifng |
| Mt2 | Mt2 | Mt2 | Mt2 |
| Crip1 | Crip1 | | |
| GzmB | GzmB | GzmB | GzmB |
| Nkg7 | Nkg7 | Nkg7 | Nkg7 |
| Ifi27l2a | | Ifi27l2a | Ifi27l2a |
| Mt3 | Mt3 | | |
| Lgals3 | | | |
| Trbc1 | Trbc1 | Trbc1 | Trbc1 |
| Batf3 | | Batf3 | Batf3 |
| S100a4 | | | |
| Ccl5 | Ccl5 | Ccl5 | Ccl5 |
| Aa467197 | | Aa467197 | Aa467197 |
| Lgals1 | Lgals1 | Lgals1 | Lgals1 |
| S100a10 | S100a10 | | |
| Anxa2 | Anxa2 | | |
| Itgb7 | | | |
| Klrd1 | Klrd1 | Klrd1 | Klrd1 |
| Isg15 | Isg15 | Isg15 | Isg15 |
| Ly6c2 | | | |
| Klf2 | Klf2 | | |
| Prf1 | Prf1 | Prf1 | Prf1 |
| Serpinb6b | Serpinb6b | | |
| Odc1 | | Odc1 | Odc1 |
| Btg1 | Btg1 | | |
| Gnai2 | | | |
| Ddt | Ddt | | |
| Bcl2a1d | Bcl2a1d | | |
| Nfkbia | Nfkbia | Nfkbia | Nfkbia |
| Cotl1 | Cotl1 | Cotl1 | Cotl1 |
| nolc1 | nolc1 | | |
| Pycard | Pycard | Pycard | Pycard |
| Rasgrp2 | | | |
| Cmtm7 | | | |
| Flna | Flna | | |
| Ctla2a | Ctla2a | | |
| Vim | Vim | Vim | Vim |
| Aurka | Aurka | | |
| Tnfrsf4 | Tnfrsf4 | | |
| Gmfg | | | |
| S1pr4 | | | |
| Ifi27 | | | |
| Chchd10 | | | |
| Zbp1 | Zbp1 | Zbp1 | Zbp1 |
| Ccr7 | Ccr7 | Ccr7 | Ccr7 |
| Id2 | Id2 | | |
| Ddx21 | Ddx21 | Ddx21 | Ddx21 |
| Chchd4 | Chchd4 | | |
| Bcl2a1b | Bcl2a1b | | |
| Top2a | Top2a | | |
| Jund | Jund | Jund | Jund |
| Npc2 | | Npc2 | Npc2 |
| Ifitm3 | Ifitm3 | Ifitm3 | Ifitm3 |
| Il2ra | Il2ra | | |
| Batf3 | Batf3 | Batf3 | Batf3 |
| Glipr2 | | | |
| Fdps | | | |
| Tspd | Tspd | Tspd | Tspd |
| Srgn | | | |
| Nefh | Nefh | | |
| Pten | Pten | Pten | Pten |
| Junb | | | |
| Cd160 | Cd160 | Cd160 | Cd160 |
| Fbxo5 | Fbxo5 | | |
| Fyb | | | |
| Cd8b1 | | | |
| Incenp | | | |
| H2ax | H2ax | | |
| Hadh | | | |
| Rras2 | | | |

| Cluster # 1 enriched motif and associated genes |
|---|
| Zf5 motif |
| Actb1 |
| Pim1 |
| Slc39a1 |
| Abhd2 |
| Alkbh5 |
| Wtap |
| Tubb5 |
| Kdelr2 |
| Pfas |
| Sephs2 |
| Gnb1 |
| Birc6 |
| Timmdc1 |
| Nup98 |
| Kmtd2 |
| Zc3hav1 |
| Actr2 |
| Polh |
| Rrn3 |
| Dpysl2 |
| Cdk6 |
| Pan3 |
| Ythdf2 |
| St13 |
| Yme1l1 |
| Arglu1 |
| Mcmbp |
| Gm2a |
| Otub1 |
| Gsk3b |
| Golt1b |
| Gtf1a1 |
